# Characterizing the demographic history and prion protein variation to infer susceptibility to chronic wasting disease in a naïve population of white-tailed deer (*Odocoileus virginianus*)

**DOI:** 10.1101/2020.10.30.362475

**Authors:** Sarah E Haworth, Larissa Nituch, Joseph M Northrup, Aaron BA Shafer

## Abstract

Assessments of the adaptive potential in natural populations are essential for understanding and predicting responses to environmental stressors like climate change and infectious disease. Species face a range of stressors in human-dominated landscapes, often with contrasting effects. White-tailed deer (deer) are expanding in the northern part of their range following decreasing winter severity and increasing forage availability. Chronic wasting disease (CWD), a prion disease affecting cervids, is likewise expanding and represents a major threat to deer and other cervids. We obtained tissue samples from free-ranging deer across their native range in Ontario, Canada which has yet to detect CWD in wild populations of cervids. We used high-throughput sequencing to assess neutral genomic variation, and variation in the PRNP gene that is partly responsible for the protein misfolding when deer contract CWD. Neutral variation revealed a high number of rare alleles and no population structure, and demographic models suggested a rapid historical population expansion. Allele frequencies of PRNP variants associated with CWD susceptibility and disease progression were evenly distributed across the landscape and consistent with deer populations not infected with CWD. We then estimated the selection coefficient of CWD, with simulations showing an observable and rapid shift in PRNP allele frequencies that coincides with the start of a novel CWD epidemic. Sustained surveillance of genomic and PRNP variation can be a useful tool for CWD-free regions where deer are managed for ecological and economic benefits.

## Introduction

Human-induced environmental change has caused widespread alterations to ecological and evolutionary processes (Harmon, Moran, & Ives, 2009; Pecl et al., 2017). Climate change is expected to be the dominant driver of wildlife population declines and has been linked to broad-scale biodiversity losses, but regional responses are often nuanced and context dependent (e.g. Taylor et al., 2017; White, Gregovich, & Levi, 2017; Hashida et al., 2020). For example, climate change in the Midwestern United States differentially favors the survival of two sympatric populations of ungulates with similar selection pressures and life history traits, but differing densities (Escobar, Moen, Craft, & VanderWaal, 2019; Weiskopf, Ledee, & Thompson, 2019). In such instances, intraspecific genetic diversity and adaptive potential is crucial for long-term population viability (Kardos & Shafer, 2018).

The emergence, spread, and persistence of infectious diseases in previously allopatric populations is facilitated by climate change and other anthropogenic activity (Price et al., 2016; Aguirre, 2017; Morand & Walther, 2020). The impacts of infectious disease on wildlife populations are of interest to managers, especially if the affected species holds economic or cultural value (Lambert et al., 2018; Weiskopf, Ledee, & Thompson, 2019). Preventing and controlling diseases in free-ranging populations can, however, be complex and costly when they are both naїve to the infectious disease and faced with climate change and other anthropogenic activities (Herrera & Nunn, 2019; Miguel et al., 2020; Samuel et al., 2020). Selective pressures are increased under these circumstances and populations are forced to respond to multiple stressors simultaneously, or potentially face local extirpation (Fischer et al., 2020).

White-tailed deer (*Odocoileus virginianus*; deer) are the most widely distributed and abundant ungulates in North America and hold significant economic and cultural value (Hewitt, 2011). The northern range of deer is primarily limited by snow but decreasing winter severity has allowed deer to expand northward beyond their historical range limits (Dawe & Boutin, 2016; Kennedy-Slaney, Bowman, Walpole, & Pond, 2018). This expansion has implications for ecosystems as, for example, deer herbivory alters long-term regional habitat characteristics and plant communities (Frerker, Sabo, & Waller, 2017; Otsu, Iijima, & Nagaike, 2019; Kroeger et al., 2020). Further, deer are an important prey species and predator populations increasing in response to deer expansion has led to greater predation and apparent interspecific competition with other ungulates (Latham et al., 2013; Barber-Meyer & Mech, 2016). Consequently, northward expansions of deer are having profound impacts to ecosystems including facilitating infectious pathogen and disease spread (Averill et al., 2018; Ferretti & Mori, 2020).

A widespread threat to deer in North America is the highly infectious and fatal neurodegenerative prion disease called chronic wasting disease (CWD). CWD is the only prion disease known to infect captive and free ranging species of cervids (family *Cervidae*) and has been reported in North and South America, Europe, and South Korea (Haley et al., 2019). With virtually no barriers to transmission and a lengthy infectious preclinical period, the local prevalence of CWD in North America has been measured to be as high as 50% and 82% in wild and captive populations, respectively (Miller et al., 2004; O’Rourke et al., 2004). Due to constraints on CWD surveillance it is likely that the distribution and prevalence of CWD in wild populations are underestimated (Escobar et al., 2020). It is clear the frequency and occurrence of CWD has increased over time, in part driven by anthropogenic activities related to hunting and wildlife farming (Osterholm et al., 2019).

Despite CWD being fatal there is inter-individual variation in susceptibility and clinical progression. Susceptibility and clinical progression are associated with non-synonymous and synonymous genetic variation in the functional prion protein gene (PRNP; Güere et al., 2020; Chafin et al., 2020). Single nucleotide polymorphisms (SNPs) at nucleotide (nt) 60, nt153, nt285, nt286, nt555, and nt676 in deer PRNP have been associated with altered CWD susceptibility or pathogenic processes (Johnson et al., 2006a; Wilson et al., 2009; Brandt et al., 2015; Brandt et al., 2018). The presence of CWD appears to affect population PRNP allele frequencies over space and time due to selection (Robinson et al., 2012); however, altered CWD susceptibility and pathogenic processes are clearly polygenic traits (Seabury et al. 2020) and disease spread is different in structured populations, which might require different wildlife management practices (Chafin et al., 2020).The efficacy of selection in the face of a novel pressure like CWD is dependent on the effective population size (*N*_e_). Common metrics to infer selection such as Tajima’s *D*, specifically measure shifts in allele frequencies across the site-frequency spectrum; however, the frequency and proportion of rare alleles is sensitive to demographic processes (Messer, Ellner, & Hairston Jr., 2016; Platt et al., 2019), and population genetics theory predicts an excess of rare alleles in expanding populations (Gillespie, 2004).

Ontario, Canada reflects the northern leading edge of deer range in eastern North America (Kennedy-Slaney, Bowman, Walpole, & Pond, 2018). The landscape of Ontario is heterogenous and environmental clines exist around the Great Lakes region. Ontario has not detected CWD in wild cervids, but CWD has been detected in farmed and captive cervids from virtually every jurisdiction bordering Ontario, with the province using a weighted surveillance approach to model CWD risk and strategically use resources for surveillance. Accordingly, at the genome-level, we predicted an excess frequency of rare neutral and PRNP variants across our study region given Ontario’s deer population is expanding (Kennedy-Slaney, Bowman, Walpole, & Pond, 2018). Based on large-scale distribution changes and recent population trends in deer (Baldwin, Desloges, & Band, 2000; Latch et al., 2009), we predicted we would observe high neutral genomic diversity (*N*_*e*_) and demographic population expansion, indicating increased gene flow and decreased population structure despite a heterogenous landscape. Since PRNP is not under selection by CWD given the region is disease free, we predicted functional variation to resemble regions most recently exposed to CWD (or still disease free). This is the first study to characterize PRNP genetic variation and population genomic structure of wild deer, while also determining the ancient and contemporary demographic and selection processes driving patterns of diversity.

## Materials and Methods

### Study area and sample collection

We sampled white-tailed deer across Ontario, Canada (Figure 1). Between 2002-2018, retropharyngeal lymph nodes were opportunistically extracted from hunter harvested deer across Ontario through the CWD surveillance program managed by the Ontario Ministry of Natural Resources and Forestry (OMNRF). Auxiliary data including year, sex, age-class, Mercator grid cell unit (GCU; 10×10 km), and wildlife management unit (WMU) were also collected with deer samples. North western and southern Ontario regions are geographically discontinuous for deer (Figure 1), samples were therefore assigned to northern Ontario or southern Ontario (Figure S1) for the purpose of analysis where sampling regions are compared. Preliminary analysis of data did not warrant separating southeastern and southwestern Ontario as per provincial management zones. Genomic DNA was extracted from deer samples using a silica-based DNA extraction kit for tissue following manufacturers protocol (Qiagen, Cat. No. 69506) and stored at −20°C. DNA quality was assessed by a spectrometer (NanoDrop 2000, Thermo Scientific) and by 2% agarose GelRed gel electrophoresis.

**Figure 1.**
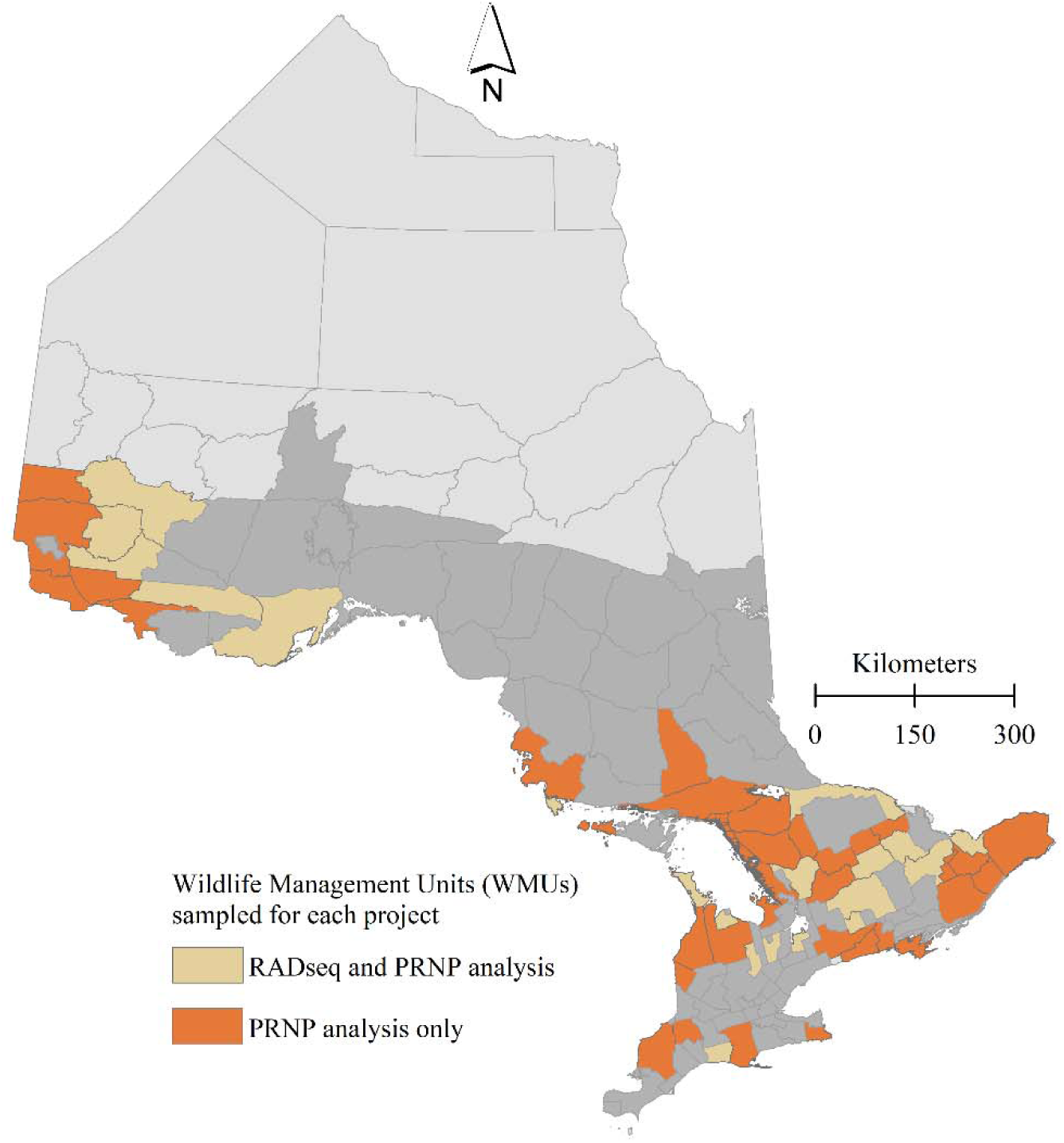
Distribution of free-ranging white-tailed deer samples obtained between 2003-2018 by the Ontario Ministry of Natural Resources and Forestry (OMNRF) that were used for the reduced representation genome analysis (n=190; cream) and the prion protein genetic analysis (n=631; orange *and* cream). The natural distribution of free-ranging white-tailed deer is shown for Ontario with a darker shade of gray.

### Library preparation for PRNP genetic analysis

A 771 base-pair (bp) region of the deer prion protein precursor (PRNP) gene was targeted and amplified using four degenerate primers (Table S1). Four replicate PCRs generated a 460 base pair Fragment 1 and a 580 base pair Fragment 2 (Table S2). In a 2:1 ratio of Fragment 1 to Fragment 2, respectively, amplified DNA was added for each individual and then indexed with standard Illumina multiplexing indices. Negative controls of UltraPure distilled water (Invitrogen, 1897011) were used for each 96-well plate. The library was purified of artifacts following manufacturers protocol for AMPure XP beads (Beckman Coulter, A63880) and validated with a TapeStation D1000 kit (Agilent, 5067-5582). The library was sequenced on an Illumina MiSeq platform at the University of Guelph Advanced Analysis Centre to generate 300 base pair (bp) pair-end reads for each sample.

### Library preparation for RADseq genomic analysis

Restriction-site associated DNA sequencing (RADseq) libraries were generated using an adapted protocol from Parchman et al. (2012) and Peterson et al. (2012) with Sbfl-HF and Msel restriction enzymes. Samples were incubated, digested overnight, and heat-inactivated in 96-well plates (Table S3). Negative controls of UltraPure distilled water (Invitrogen, 1897011) were used for each 96-well plate. Restriction digested DNA was combined with 7 ul of ligation mixture and 3 ul of one of the 24 available Sbfl adapters (1.0 uM). Adapters were ligated at 16°C for 3 hours. DNA fragments were purified of artifacts following manufacturers protocol for AMPure XP beads (Beckman Coulter, A63880). Adapter-ligated fragments were amplified in four separate 10 ul reactions that incorporated barcodes. Reaction conditions and primers are shown for ligation mixture and PCR in Table S4 and Table S5, respectively. Samples were pooled and purification was performed following manufacturers protocol for QIAquick PCR Purification kit (Qiagen, 28106) for a final elution to 42 ul. Size selection between 450 bp to 700 bp was performed on 80 ul replicates of purified library and gel purification was performed following manufacturers protocol for QIAquick Gel Extraction kit (Qiagen, 28706) for a final elution to 60 ul. The purified final library was validated with a TapeStation D1000 kit (Agilent, 5067-5582). The libraries were sequenced at The Centre for Applied Genomics (TCAG) in The Hospital for Sick Children (SickKids) on an Illumina HiSeq 2500 to produce 2×126 base pair paired end reads.

### Bioinformatic pipeline and data analysis I – PRNP gene

The quality of reads was assessed using FastQC (Babraham Institute; v0.11.8). Samples were excluded if at least one file in the pair-end files for a sample was less than 1kB in size or failed to pass quality standards. A novel command line-based pipeline was developed to assemble and genotype PRNP (accessible at https://gitlab.com/WiDGeT_TrentU). Our workflow integrated Pullseq v1.0.2, BWA v0.7.17, SAMtools; v1.9, BCFtools v1.9, and VCFtools v0.1.16-15. Briefly, for each sample, the pipeline extracted relevant reads based on the presence of primer sequence, mapped the extracted reads to a 771 bp PRNP gene reference sequence. We generated a consensus sequence and called single nucleotide polymorphisms (SNPs). SNP calls were limited to positions where there was a minimum read depth of 30 and mapping quality score of at least 30. Sanger sequencing of a subset of samples and their a priori called variants were used to validate the bioinformatic pipeline.

The presence of asparagine (N) at aa138 (nt413A) indicates amplification of the pseudogene (Brandt et al., 2015); we therefore filtered out all sequences with this site. SNPs with a total frequency of occurrence of 1% or less were excluded from the analysis. A two-sided Fisher’s Exact Test was conducted on minor allele counts from either northern or southern sampling regions at four well studied positions in the deer PRNP gene associated with CWD: nucleotide (nt) 60, nt285, nt286, and nt676. Synonymous and non-synonymous sites were identified using MEGA X v10.0.5. Haplotypes were estimated from unphased sequences with PHASE v2.1.1 using a Markov chain Monte Carlo (MCMC) sampling approach with a minimum of 100,000 steps, with a discarded burn-in of 10,000, and samples were drawn every 100 MCMC steps. Five repetitions were performed to verify consistent frequencies of haplotype assignment (Brandt et al., 2018). Haplotypes with a frequency of less than 1% were removed. The genotype, frequencies, and estimated standard deviations of the remaining haplotypes were analyzed as a 2×2 contingency table by sampling region.

### Bioinformatic pipeline and data analysis II – RADseq

Fastq files were demultiplexed using process_radtags within the Stacks v2.3 module. Parameters within process_radtags included the removal of any read with an uncalled base and the discarding of reads with low quality scores. The demultiplexed sample files were aligned against the deer genome (Genome Accession JAAVWD000000000) using BWA with samtools used to sort, merge and compress BAM files. The referenced-based approaches on gstacks and populations program within STACKs produced a variant call format (VCF) file with the restrictions that the minimum percentage of individuals in a population required to process a locus for that population was 90%. The VCF was filtered using VCFtools to only include reads with a minimum read depth of 20. Population statistics, including FIS, observed and estimated homozygosity, and nucleotide diversity were calculated using the *populations* module. To analyze sources of variation, we generated a principal component analysis (PCA) using the R v3.6.1 package adegenet v2.1.3. A linear regression was run on principal component (PC) 1 and PC2 scores against latitude and longitude. We estimated FST between north and south using StAMPP v1.6.1. Population structure was detected using successive K-means clustering and a discriminant analysis of principal components (DAPC) available in adegenet (Jombart, Devillard, & Balloux, 2010).

### Demographic Analysis and Estimate of Effective Population Size

The final VCF was converted into 1D site frequency spectrum (SFS) for all of Ontario and northern vs and southern designations, respectively, using vcf2dadi.py with projections for the SFS estimated in easySFS. We applied a diffusion-based approach to demographic inference through the Diffusion Approximation for Demographic Inference (δaδi) tool by Gutenkunst et al., (2009). Nine 1D models were assessed for Ontario as a single population. The optimum model was selected as the lowest optimized log-likelihood of all successfully run models. δaδi was also used to estimate the following summary statistics for the province: Watterson Theta (θ), Tajima’s *D* and the number of segregating sites. Using the mutation rate (*μ*) per site per generation of a closely related species (*Rangifer tarandus* from Chen et al., 2019), total number of sites (L), and the parameters estimated in the optimum model selected from δaδi, we estimated the ancestral effective population size as *N*_*a*_ = θ / 4*μ*L.

### Estimation of selection on PRNP and allele frequency projections for a naїve population

We estimated the selection coefficient (*s*) at two sites (nt285 and nt286) using the approach of Thompson et al. (2019) which accounts for number of generations and different expression modes (i.e. recessive, dominant, codominant). Strength of selection against the less resistant phenotype [i.e., the homozygous common allele (*s*_AA_)] can be estimated by calculating values of sAA that explained the estimated change in allele frequencies between positive and negative animals from the same region. Here we used starting (negative CWD) allele frequencies from Wilson et. al. (2009) and Kelly et al. (2009) and estimated *s* over *n* generations the equation:

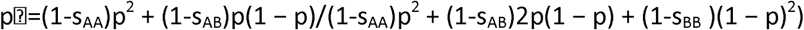

taken from Charlesworth and Charlesworth (2010). To account for uncertainty in time and allele frequency estimates we ran 100 iterations with *n* ranging from 2-25 generations (∼4-50 deer years) and positive and negative allele frequencies (±1%) for those reported from both Wilson et. al. (2009) and Kelly et al. (2009). Then using our estimated allele frequencies for Ontario, we projected each allele frequencies into the next 25 generations using our estimated selection coefficients, and the estimated *s* coefficients of 0.0103 and 0.074 from Robinson et al. (2012). Calculations were conducted under three relative fitness scenarios as per Thompson et al. (2019).

## Results

### PRNP Genetic analysis

A total of 631 Ontario deer samples were included in the PRNP genetic analysis (Figure 1). Nineteen SNPs were detected after filtering (Table 1), with 8 being non-synonymous substitutions. Six of the detected variants in the PRNP gene have been linked to CWD susceptibility or clinical progression, these include nt60, nt153, nt285, nt286, nt555, and nt676. A two-sided Fisher’s Exact test on the major and minor allele counts at the four important, arguably the most studied, CWD-linked loci (nt60, nt285, nt286, and nt676) conducted between Northern and Southern Ontario indicated that there was only a difference in frequency (p < 0.05) at nt676 (Table S6).

**Table 1.** Allele and amino acids frequencies for reference/alternate alleles at Ontario white-tailed deer (n=631) prion protein gene.

| Locus | Allele | Codon | Amino acid | Major frequency | Minor frequency |
| --- | --- | --- | --- | --- | --- |
| 60 | C/T | 20 | D/- | 0.9794 | 0.0206 |
| 136† | A/G | 46 | S/G | 0.9873 | 0.0127 |
| 153 | C/T | 51 | R/- | 0.8605 | 0.1395 |
| 168† | A/G | 56 | G/- | 0.9635 | 0.0365 |
| 174† | T/G | 58 | G/- | 0.9842 | 0.0158 |
| 177† | C/G | 59 | G/- | 0.9873 | 0.0127 |
| 178† | T/G | 60 | W/G | 0.9160 | 0.0840 |
| 192† | T/G | 64 | H/Q | 0.8653 | 0.1347 |
| 195† | A/G,T | 65 | G/- | 0.7147 | 0.2853 |
| 198† | T/G | 66 | G/- | 0.7021 | 0.2979 |
| 285 | A/C | 95 | Q/H | 0.9699 | 0.0301 |
| 286 | G/A | 96 | G/S | 0.6609 | 0.3391 |
| 324 | A/G | 108 | P/- | 0.9794 | 0.0206 |
| 365† | G/T | 122 | G/V | 0.9651 | 0.0349 |
| 378 | G/A | 126 | G/- | 0.9651 | 0.0349 |
| 417 | G/A | 139 | R/- | 0.1696 | 0.8304 |
| 548† | T/A | 183 | V/D | 0.9778 | 0.0222 |
| 555 | C/T | 185 | I/- | 0.4120 | 0.5880 |
| 676 | C/A | 226 | Q/K | 0.9620 | 0.0380 |

† indicates novel positions.

There were 102 unique haplotypes with a count of at least one, with 12 haplotypes having a frequency greater than 1% (Figure 2; Table S7). The two most common haplotypes, Haplotype 3 (*f*=0.23) and Haplotype 1 (*f*=0.12) did not include any non-synonymous substitutions. Haplotype A (*f*=0.30) and Haplotype B (*f*=0.25) reported by Brandt et al (2015, 2018) from northern Illinois were also detected as Haplotype 16 (*f*=0.09) and Haplotype 7 (*f*=0.05), respectively. The same Haplotype A (*f*=0.15) and Haplotype B (*f*=0.23) were reported by Chafin et al., 2020 in deer from Arkansas, USA.

**Figure 2.**
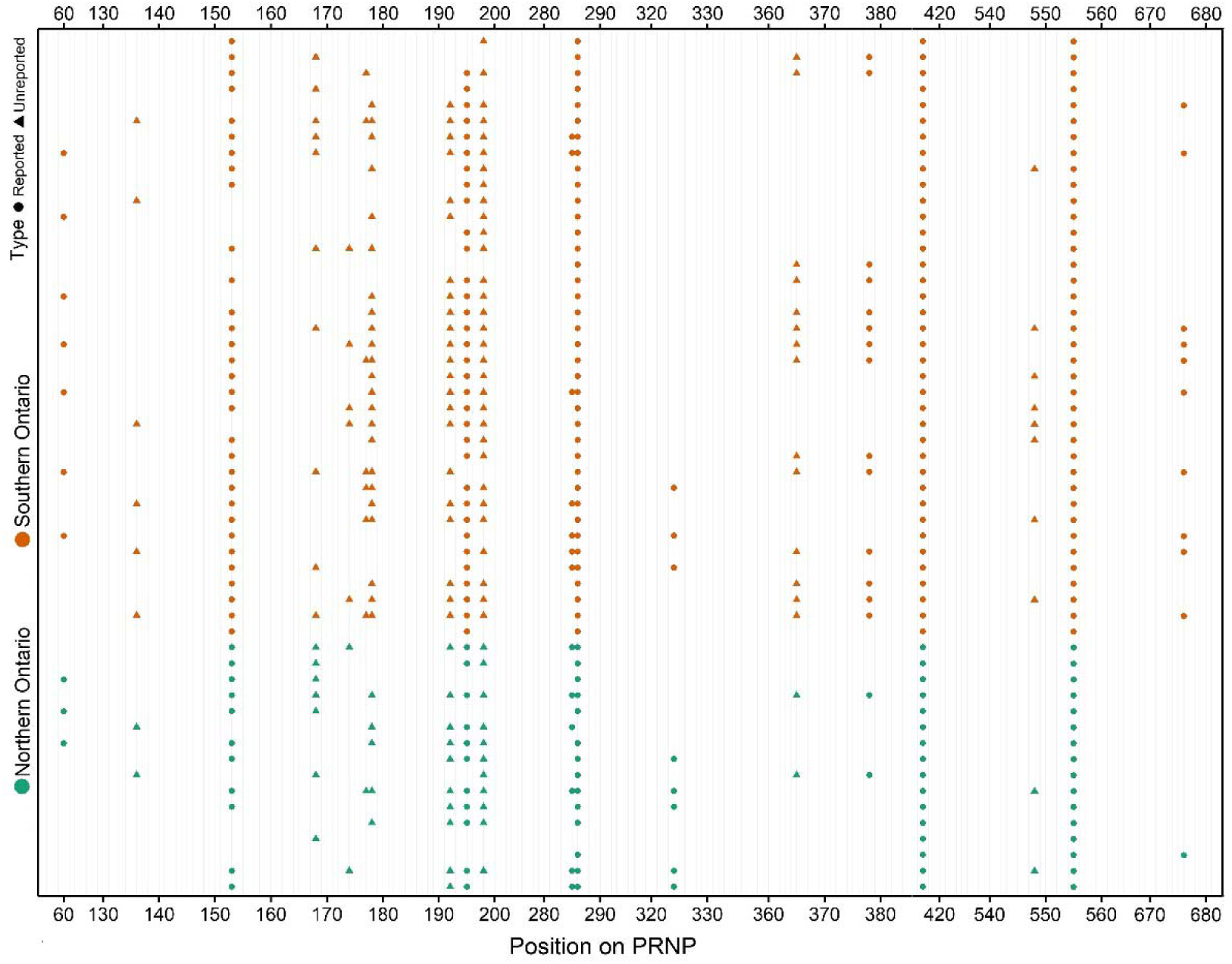
A 771 bp region of the white-tailed deer prion protein gene was analyzed from free-ranging white-tailed deer in Ontario, Canada (n=631). The overlayed genotypes across 19 variable loci were organized by broad management in Ontario and are shown. Circles indicate loci previously described as variable in the white-tailed deer prion protein gene. Triangles indicate novel variable loci in the variable loci in the white-tailed deer prion protein gene.

### RADseq Genomic analysis

A total of 235 Ontario deer samples were sequenced in ddRADseq libraries. Following quality control and quality assessment, including FastQC and line counts of demultiplexed files, 190 samples remained for downstream analysis (Figure 1). Estimated population diversity statistics are summarized in Table 2. The PCA clearly separated northern and southern Ontario along PC1 (Figure 3). A linear regression revealed that PC1 was strongly associated with longitude (β = –0.79; adjusted R^2^ = 0.77; p-value < 0.01). However, population structure between northern and southern Ontario was weak (FST=0.02). A BIC based on the K-cluster analysis also indicated that the most optimum number of clusters was 1.

**Table 2.**
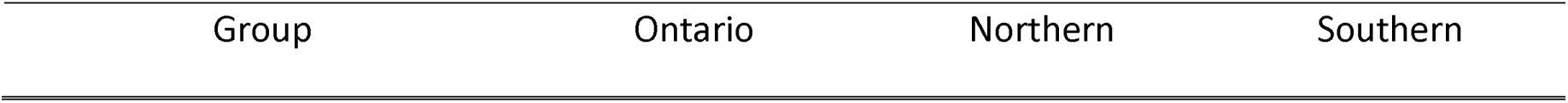
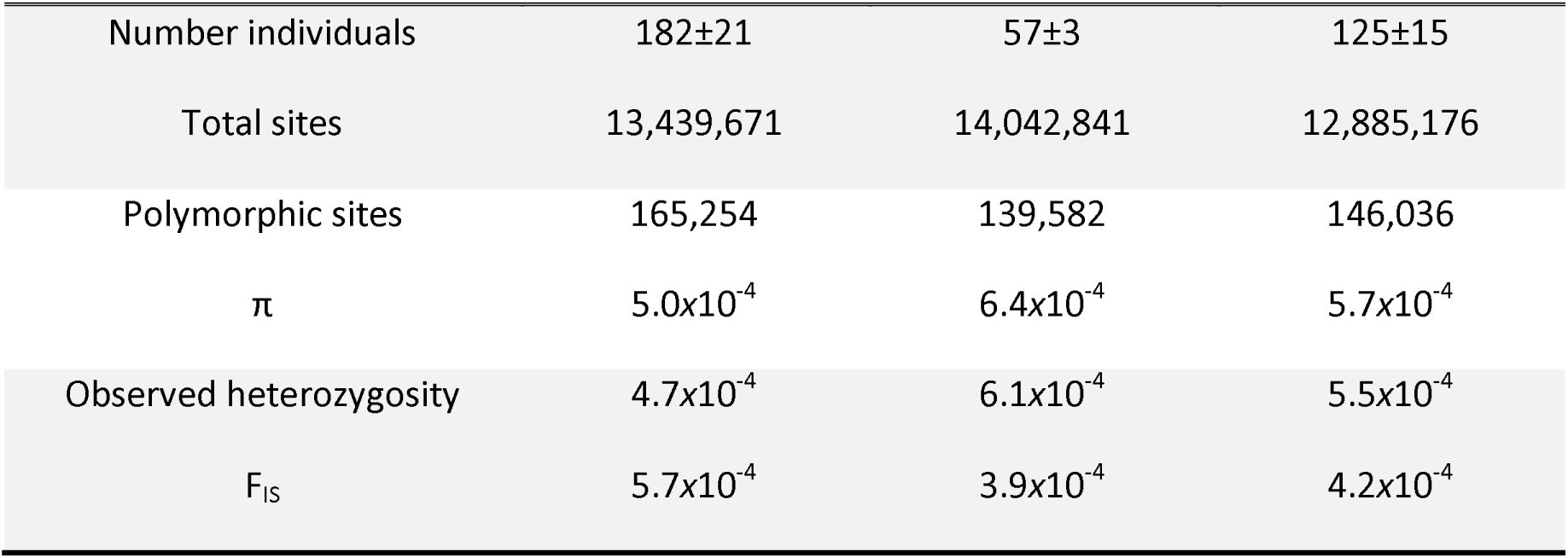
Genome-wide population summary statistics including breakdown of sites, number of individuals, nucleotide diversity estimate (π), individual genetic variance (I) relative to the subpopulation genetic variance (F_IS_), and the observed heterozygosity of white-tailed deer in Ontario.

| Group | Ontario | Northern | Southern |
| --- | --- | --- | --- |
| Number individuals | 182±21 | 57±3 | 125±15 |
| Total sites | 13,439,671 | 14,042,841 | 12,885,176 |
| Polymorphic sites | 165,254 | 139,582 | 146,036 |
| $\pi$ | $5.0 \times 10^{-4}$ | $6.4 \times 10^{-4}$ | $5.7 \times 10^{-4}$ |
| Observed heterozygosity | $4.7 \times 10^{-4}$ | $6.1 \times 10^{-4}$ | $5.5 \times 10^{-4}$ |
| $F_{IS}$ | $5.7 \times 10^{-4}$ | $3.9 \times 10^{-4}$ | $4.2 \times 10^{-4}$ |

**Figure 3.**
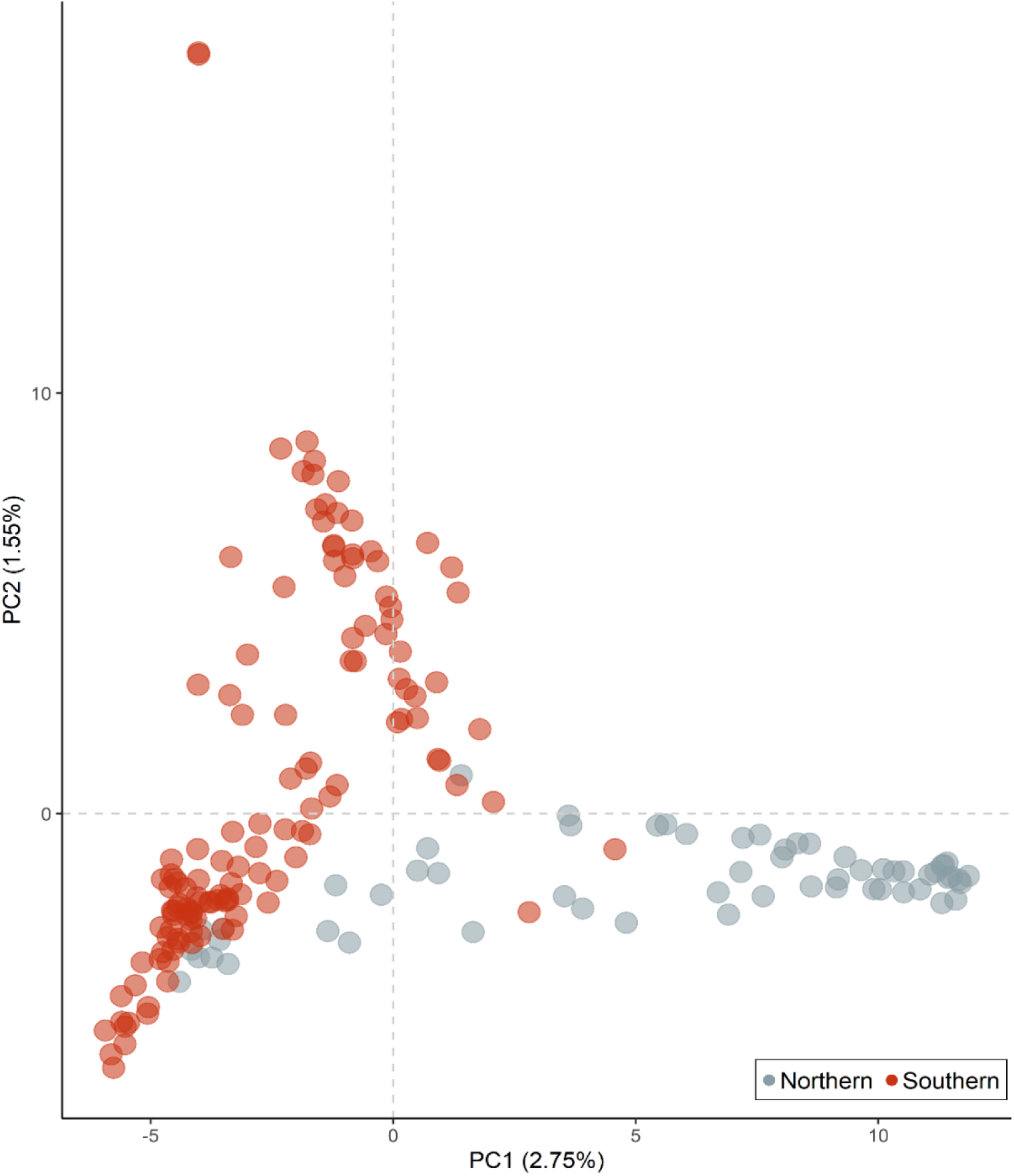
A plot of PC1 and PC2 from the principal component analysis (PCA) on the reduced representation white-tailed deer genome. PC1 and PC2 were able to explain a total of 4.3% of the genomic variation observed. Gray circles represent scores from samples in northern Ontario. Orange circles represent scores from samples in southern Ontario.

The optimum 1D demographic model for Ontario was the BOTTLEGROWTH model, which models an instantaneous size change followed by exponential change (Table S8; Figure 4). From the 1D site frequency spectrum, the mean Tajima’s *D* was estimated to be −2.126 consistent with a population expansion after a bottleneck. Using the calculated θ, we estimated *N*_*a*_ to be ∼20,000: this would place the timing of the population change (Tc) measured in 2N_a_ generations around the onset of the last glacial maximum (Figure 4).

**Figure 4.**
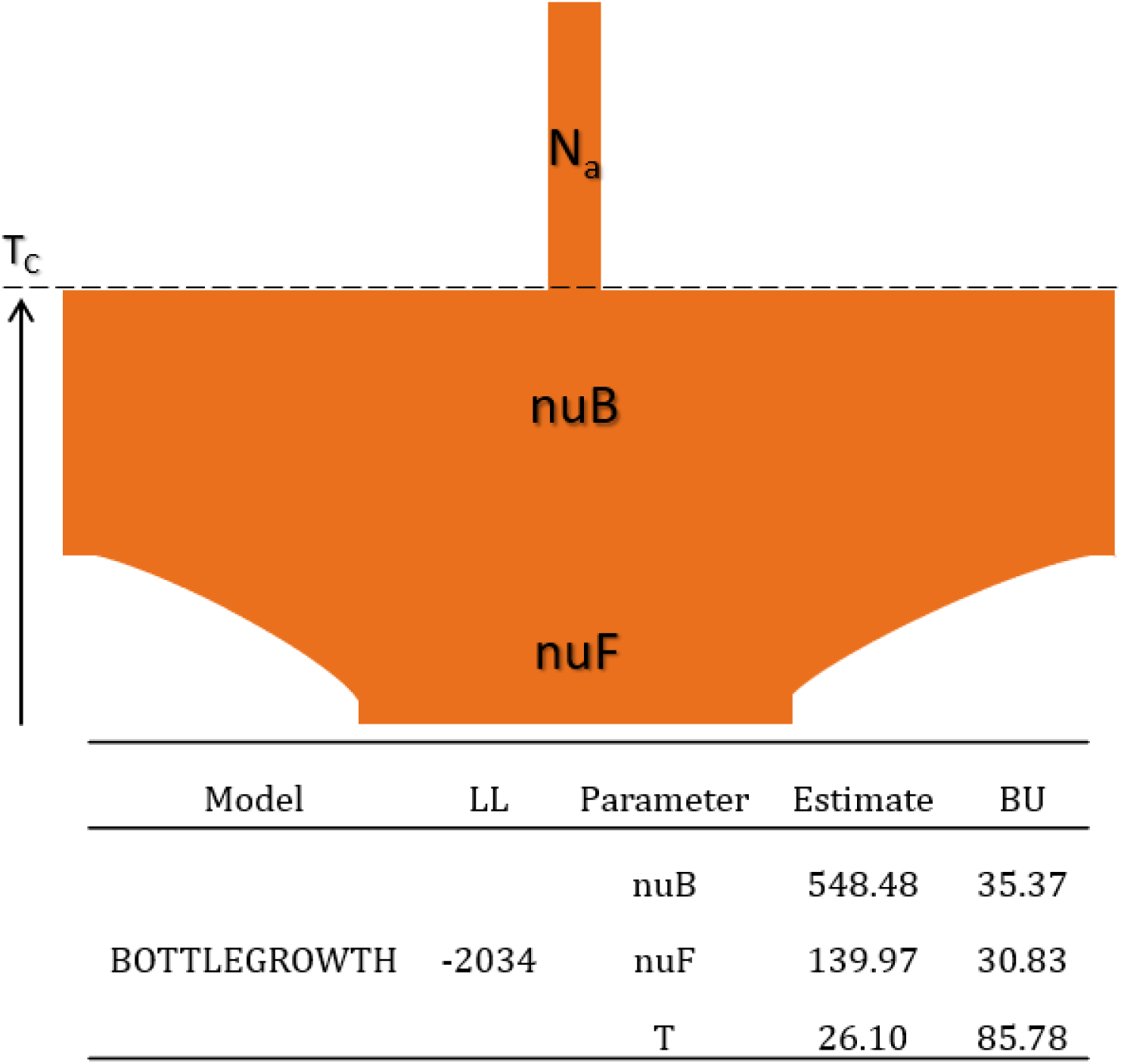
Demographic parameter estimates from δaδi for the optimal 1D model for white-tailed deer in Ontario. Parameter estimates are the ancient population size (N_a_), the ratio of population size after instantaneous change to ancient population size (nuB); the ratio of contemporary to ancient population size (nuF); and the time in the past at which instantaneous change happened and growth began (TC; in units of 2*N_a_ generations). Included are the optimized log-likelihood (LL) and bootstrap uncertainties (BU).

### PRNP selection and projection

We estimated the selection coefficients to be 0.08 (±0.06) and 0.11 (±0.07) for nt285 and nt286 under a dominance model. All simulated projections with our *s* values and the upper reported value of Robinson et al (2012) showed a rapid shift in allele frequencies (Figure 5); the majority of simulated trajectories did not overlap with the low *s* coefficient of 0.01 previously reported by Robinson et al (2012).

**Figure 5.**
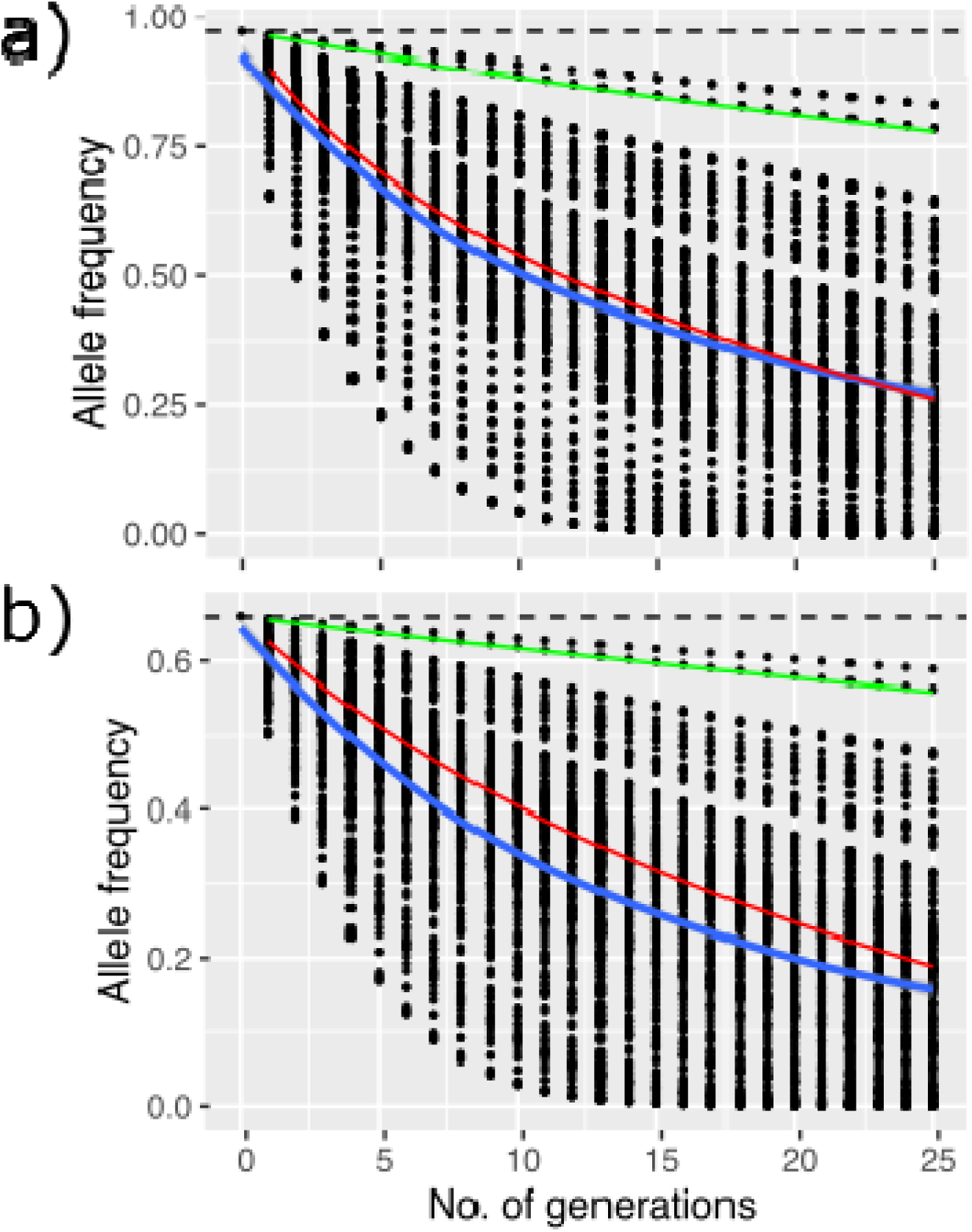
Simulated allele frequency projections for nucleotide positions 285 (a) and 286 (b) in the PRNP gene under selection. Green and red lines are the selection coefficients provided by Robinson et al (2012); while the black dots are those from our simulations, with blue line representing the mean selection coefficient.

## Discussion

White-tailed deer in North America are intensively managed for hunter harvest and are expanding their range northward due to climate change. The frequency and occurrence of CWD infection in captive and free-ranging deer is likewise increasing (Osterholm et al., 2019; Rivera, Brandt, Novakofski, & Mateus-Pinilla, 2019), but transmission and spread in free-ranging populations are still poorly understood (Potapov, Merrill, Pybus, & Lewis, 2016). We observed that the frequency of PRNP alleles in our naїve population differed from areas currently infected with CWD or where CWD is endemic, including western Canada (Table S9; Kelly et al., 2008; Wilson et al., 2009; Brandt et al., 2015; Brandt et al., 2018; Chafin et al., 2020).

The emergence, transmission, and persistence of highly infectious diseases in healthy populations are often facilitated by climate change and exacerbated in areas with intense anthropogenic activity (McKnight et al., 2017; Rizzoli et al., 2019). The introduction and re-introduction of infectious diseases often results in rapid local population declines, reducing species’ adaptive potential and generating substantial economic losses (Belant & Deese, 2010; Escobar, Moen, Craft, & VanderWaal, 2019). In Ontario deer, however, we observed an excess of rare alleles with no evidence of strong population structure, including in southern Ontario where an environmental cline exists around lake systems. An excess of rare alleles and high *N*_*e*_ and absent population structure might provide the means for effective adaptation to selective pressures, including climate change, infectious disease, and human activity.

### Linking Population Demographics to Functional genes

We assessed contemporary and historical gene flow by examining neutral genomic variation across tens of thousands of loci from deer across the province on Ontario. The data were consistent with a large expanding population with a high level of genomic diversity and an excess of rare alleles. These genomic patterns are also consistent with population simulations of responses to climate change (Kennedy-Slaney, Bowman, Walpole, & Pond, 2018), and offers some clues as to the potential adaptive response were CWD to arrive. Specifically, two important features of the genomic data are noteworthy: the high number of a rare alleles and the limited population structure across Ontario.

Demographic processes and life history strategies influence the proportion of rare alleles, which are important to the adaptive process but are sensitive to recent evolutionary processes (Peart et al. 2020). Evidence suggests the accumulation of rare alleles is independent of taxa, but adaptation appears limited in low-diversity taxa (e.g., primates; Rousselle et al., 2020). The deer genomic data evaluated here are consistent with high adaptive potential; specifically, we calculated an *N*_*a*_ to be ∼20,000, with the expansion estimated to be 100X suggestive of a large species-wide *N*_*e*_ (derived either from π in Table 2 or the demographic model in Figure 4).

Studies of populations infected with CWD often demonstrate a lack of allelic diversity at the PRNP gene, which is thought to be due to CWD-driven selection positively selecting for functionally relevant alleles (Haley et al., 2019). We found a high frequency of rare PRNP alleles in our naїve population which differs from areas currently infected or where CWD is endemic, including western Canada (Johnson et al., 2006a; Kelly et al., 2008; Wilson et al., 2009; Brandt et al., 2015; Brandt et al., 2018). Furthermore, infectious diseases can cause population fragmentation (i.e., structure), demographic changes, and genetic isolation (McKnight et al., 2017). A pre-infection presence of a large proportion of rare alleles in both functional and neutral genes, as observed in Ontario, supports surveillance data showing no CWD, which might bode well for a long-term adaptive response to CWD infection and other stressors. Importantly, even low selection coefficients should alter allele frequencies in a detectable manner (Figure 5). However, it is not clear how this standing genetic variation compares to infected populations prior to the arrival of CWD, creating some uncertainty in the potential adaptive response, recognizing that loci outside PRNP are also involved (Seabury et al. 2020).

We observed no evidence of strong population structure despite the heterogeneous landscape observed in Ontario, but there is a clear latitudinal cline in allele frequencies. This suggests random mating is largely occurring a regional levels in spite of substantial environmental changes including intense agricultural practices, substantial urbanization of the landscape, and climate change (Schulte et al., 2007; Walter et al., 2009; Patton, Russell, Windmuller-Campione, & Frelich, 2018). We might therefore expect that the spread of advantageous rare alleles under selection may not be limited in the province. Conversely, we might also expect the homogenous population to facilitate the spread of CWD if anthropogenic activity were to introduce the disease into the focal population (Escobar et al., 2020). The movement of infected wildlife might also pose a potential risk to human health at the wildlife-human-livestock interface as CWD can cross species barriers (Igel-Egalon et al., 2020). Sustained monitoring across CWD-free regions where deer are managed for sustainability or where food security is threatened should continue, but consider population-level responses to climate change (i.e. Kennedy-Slaney, Bowman, Walpole, & Pond, 2018) while integrating genetic information beyond PRNP allele frequencies.

## Conclusion

We gauged the adaptive potential of CWD naїve deer in Ontario, Canada by assessing functional and neutral genetic diversity. The genome-wide data were consistent with a large expanding population with a high neutral genomic diversity, no population structure, and an excess of rare alleles, which is also consistent with non-genetic population and climate models (Kennedy-Slaney, Bowman, Walpole, & Pond, 2018). We suggest these patterns might favour a novel adaptive response to the arrival of CWD. Voluntary hunter-harvest based surveillance for CWD will likely be able to detect the introduction of CWD in a naїve population; however, we expect that there will be a lag in detection when prevalence is low since low densities likely limit transmission (Gagnier, Laurion, & DeNicola, 2020). Unfortunately, a lag in detection will likely permit the establishment of CWD and, over time, eradication becomes nearly impossible and costs are high (Mysterud & Rolandsen, 2018). Sustained temporal monitoring of variation in PRNP across CWD-free regions could be a detection tool as our simulations suggest detectable shifts should occur with the arrival of the disease. By combining demographic patterns and genotypes with current risk models, managers could improve risk-based detection efforts and facilitate a more effective resource deployment plan as the disease alters the population.

## Acknowledgements

Natural Sciences and Engineering Research Council of Canada (NSERC) Discovery Grants (JMN - RGPIN-2019-04330, ABAS RGPIN-2017-03934), NSERC CGS (SH), Canadian Foundation for Innovation (JELF #66905), Compute Canada RRG (gme-665-AB), and the Ontario Animal Health Network provided support for the project. We thank Cathy Cullingham for comments, and Anh Dao for providing samples.

## Data Accessibility

DNA sequences: SRA Accessions PRJNA565222 and SUB8436966

PRNP and RADseq analysis pipelines: https://gitlab.com/WiDGeT_TrentU/

## Supplementary Materials

**Table S1.**
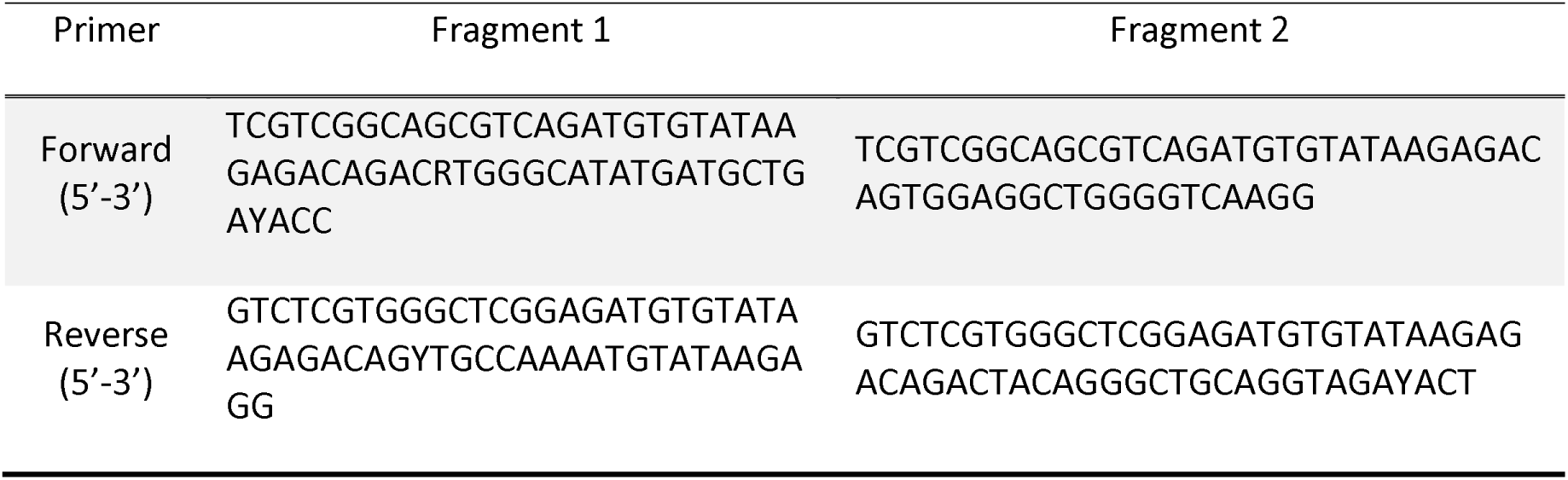
Four degenerate primers used to amplify the functional white-tailed deer prion protein gene region as two overlapping fragments are shown. Fragment 1 is 460 base-pairs in length and Fragment 2 is 590 base-pairs in length.

**Table S2.**
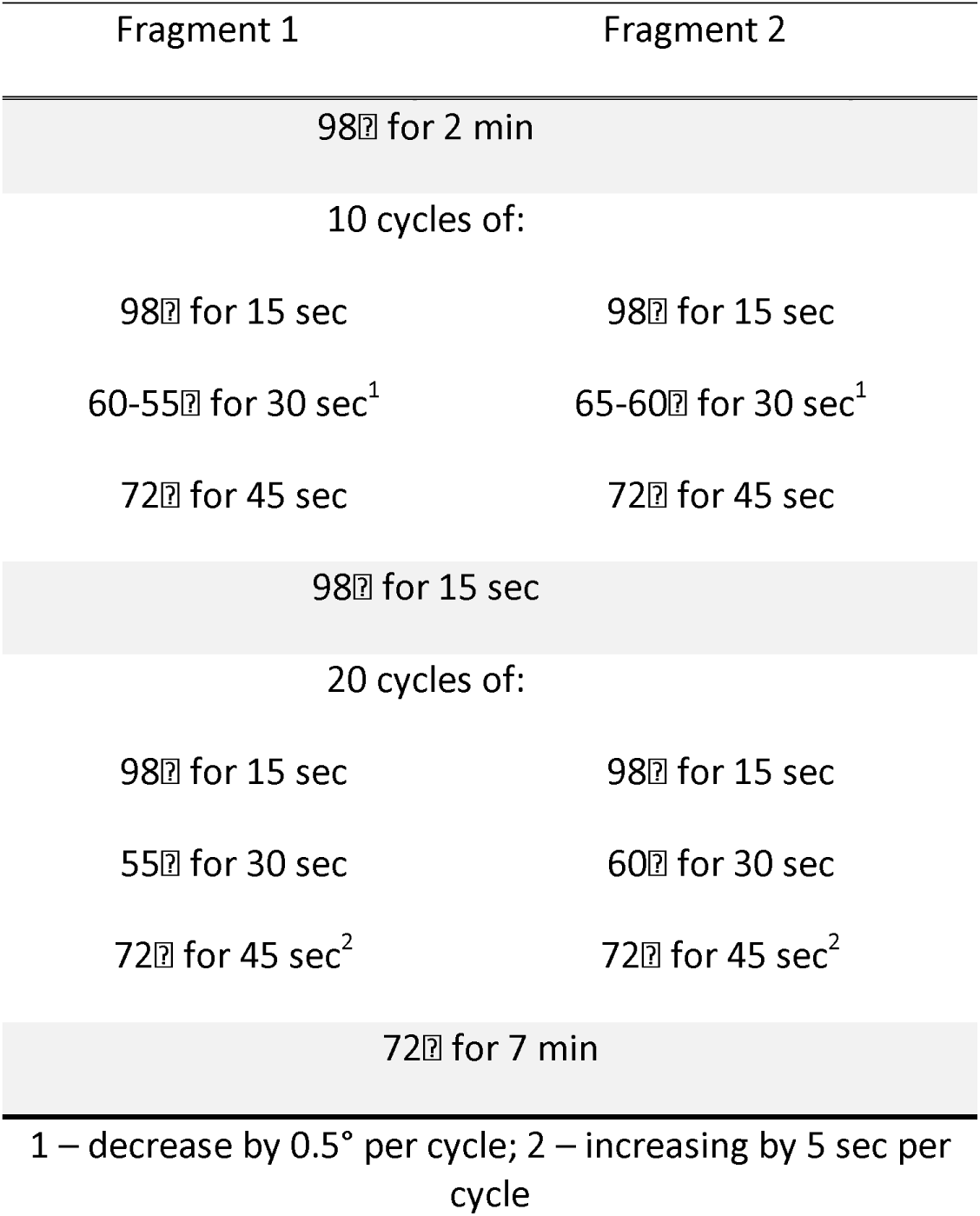
Thermal cycling conditions for PRNP PCR amplification for Fragment 1 and Fragment

| Fragment 1 | Fragment 2 |
| --- | --- |
| 98°C for 2 min |  |
| 10 cycles of: |  |
| 98°C for 15 sec | 98°C for 15 sec |
| 60-55°C for 30 sec<sup>1</sup> | 65-60°C for 30 sec<sup>1</sup> |
| 72°C for 45 sec | 72°C for 45 sec |
| 98°C for 15 sec |  |
| 20 cycles of: |  |
| 98°C for 15 sec | 98°C for 15 sec |
| 55°C for 30 sec | 60°C for 30 sec |
| 72°C for 45 sec<sup>2</sup> | 72°C for 45 sec<sup>2</sup> |
| 72°C for 7 min |  |
| 1 – decrease by 0.5° per cycle; 2 – increasing by 5 sec per cycle |  |

**Table S3.**
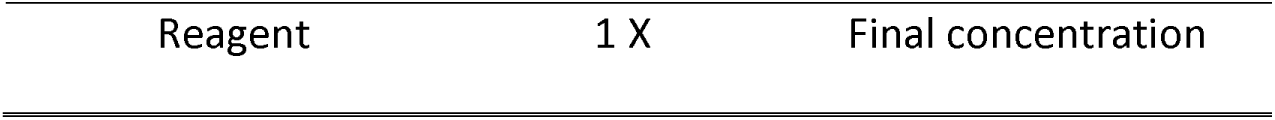
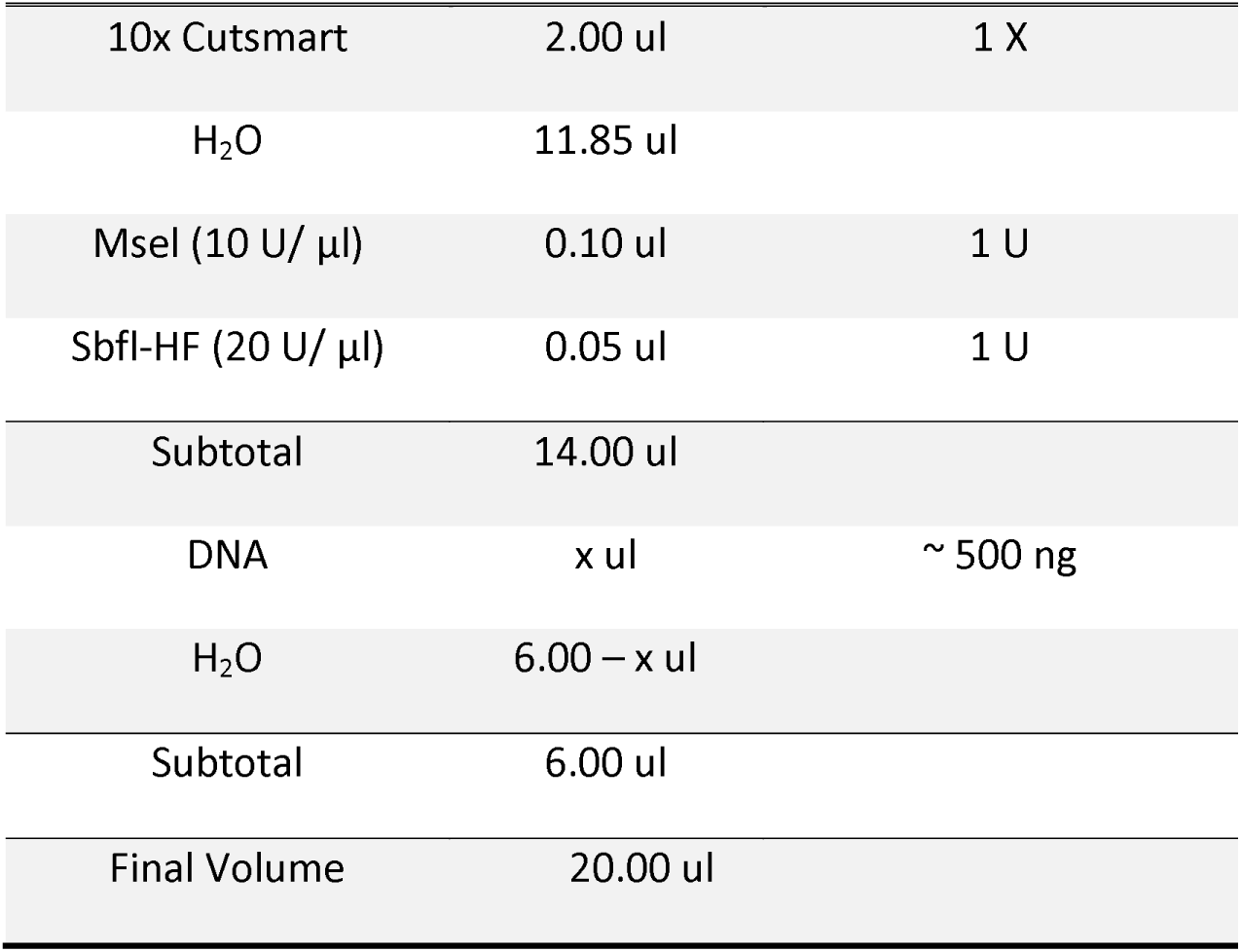
Mixture for double restriction enzyme digestion. Samples and reaction mixture were incubated at 37°C for 3 hours and then 25°C overnight on a thermal cycler with a heated lid. Samples and reaction mixture were head-inactivated at 85°C for 30 minutes.

**Table S4.**
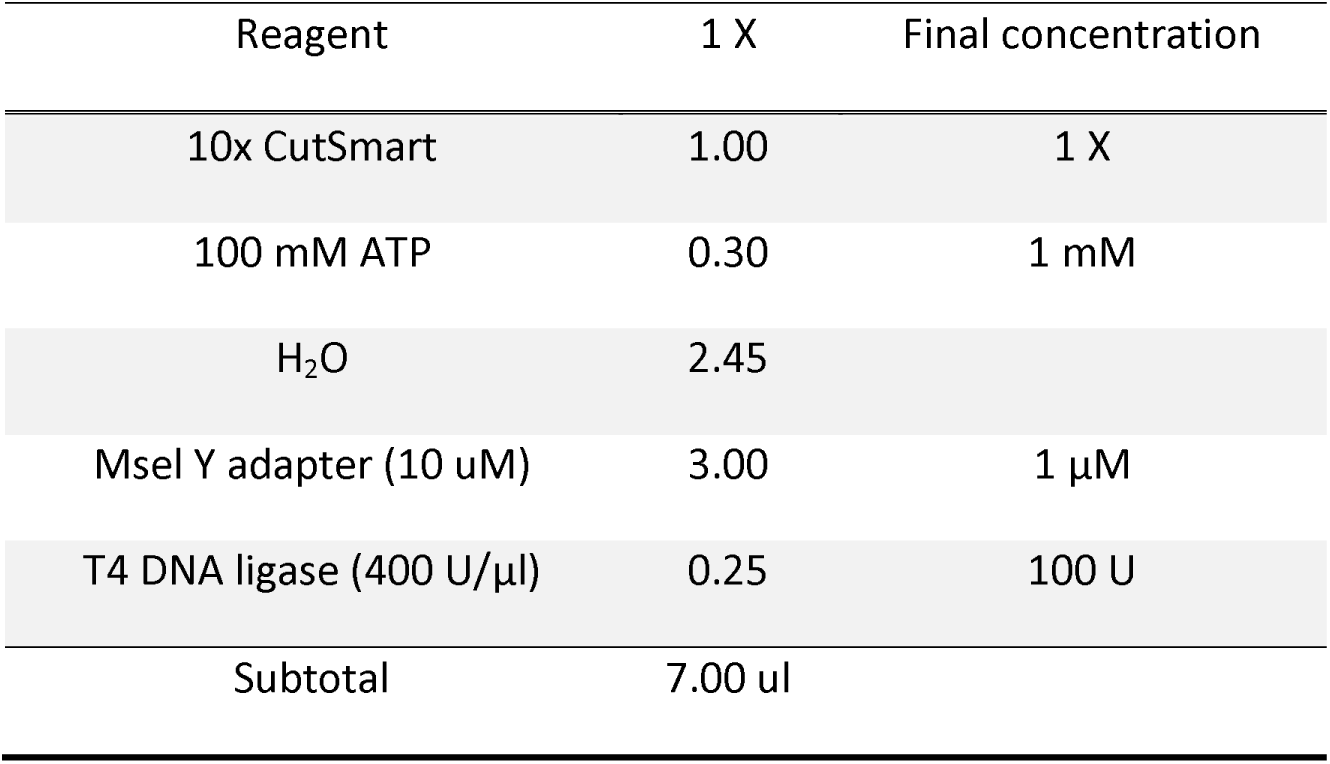
Mixture for adapter ligation. The digested fragments were combined with 7 ul of the adapter ligation mixture and 3 ul of a unique barcoded Sbfl adapter (1.0 uM). The ligation on ∼ 30 ul reaction mixture was performed in 16°C for 3 hours

| Reagent | 1 X | Final concentration |
| --- | --- | --- |
| 10x CutSmart | 1.00 | 1 X |
| 100 mM ATP | 0.30 | 1 mM |
| H<sub>2</sub>O | 2.45 |  |
| MseI Y adapter (10 uM) | 3.00 | 1 μM |
| T4 DNA ligase (400 U/μl) | 0.25 | 100 U |
| Subtotal | 7.00 ul |  |

**Table S5.**
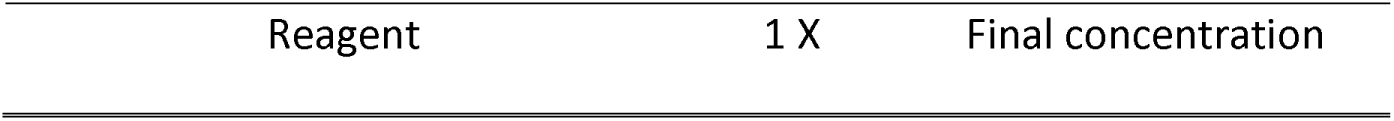
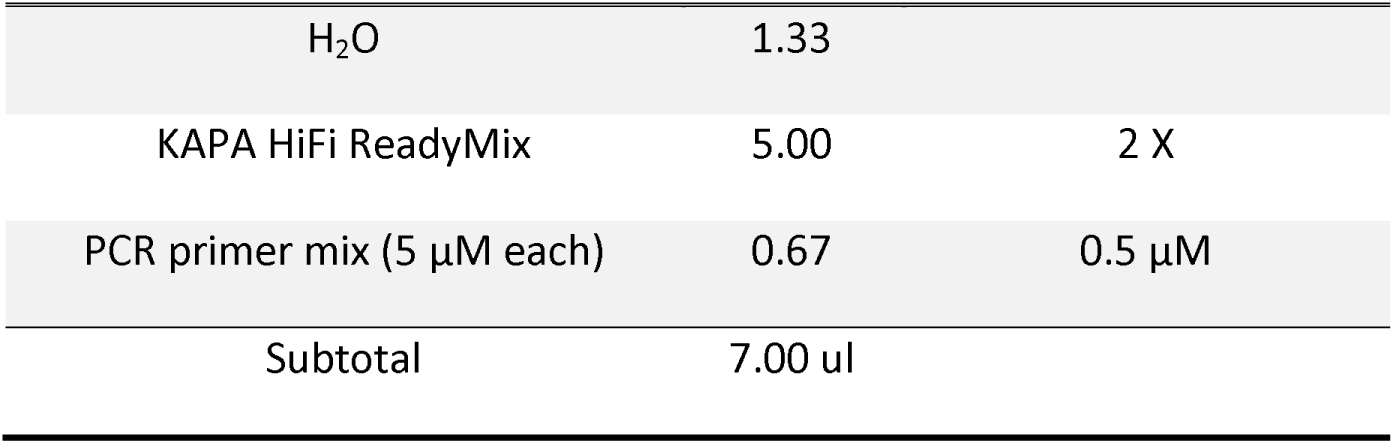
Illumina PCR mixture. The purified restriction-ligation DNA (3 ul) was combined with 7 ul of PCR mixture and a PCR was performed on 10 ul. Four replicated per sample was performed. The thermal cycler profile for this PCR was 98°C for 30 seconds; 20 cycles of 98°C for 20 seconds, 60°C for 30 seconds, 72°C for 45 seconds; and a final extension at 72°C for 5 minutes.

| Reagent | 1 X | Final concentration |
| --- | --- | --- |
| H<sub>2</sub>O | 1.33 |  |
| KAPA HiFi ReadyMix | 5.00 | 2 X |
| PCR primer mix (5 µM each) | 0.67 | 0.5 µM |
| Subtotal | 7.00 ul |  |

**Table S6.**
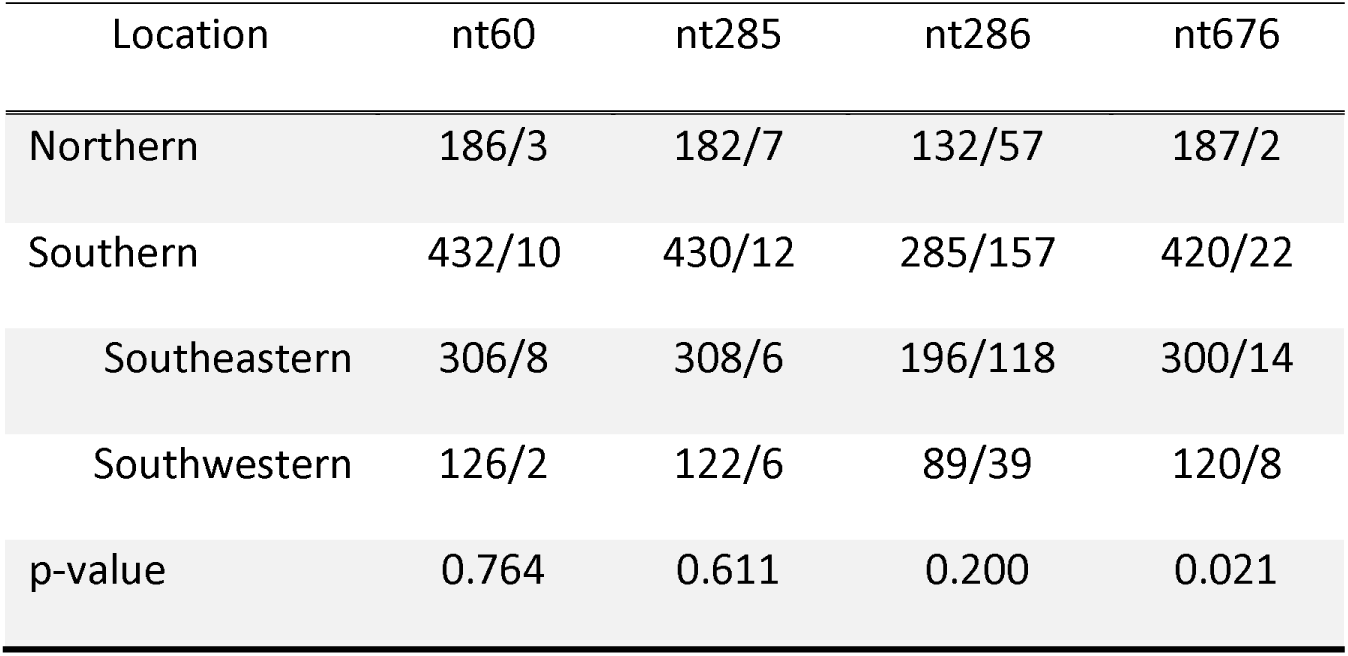
The major/minor allele counts for four nucleotide (nt) positions of variation in the white-tailed deer prion protein gene that are linked to reduced susceptibility or reduced clinical progression of chronic wasting disease for Northern and Southern Ontario are shown. A two-sided Fisher’s Exact test was conducted on the major and minor allele counts at the four chronic wasting disease-linked nucleotide positions. A p-value less than 0.05 was considered significant and indicated that there were significant differences between the groups.

**Table S7.**
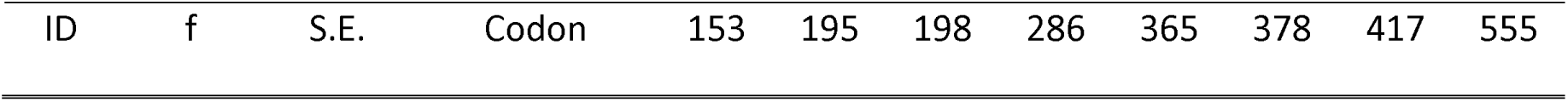
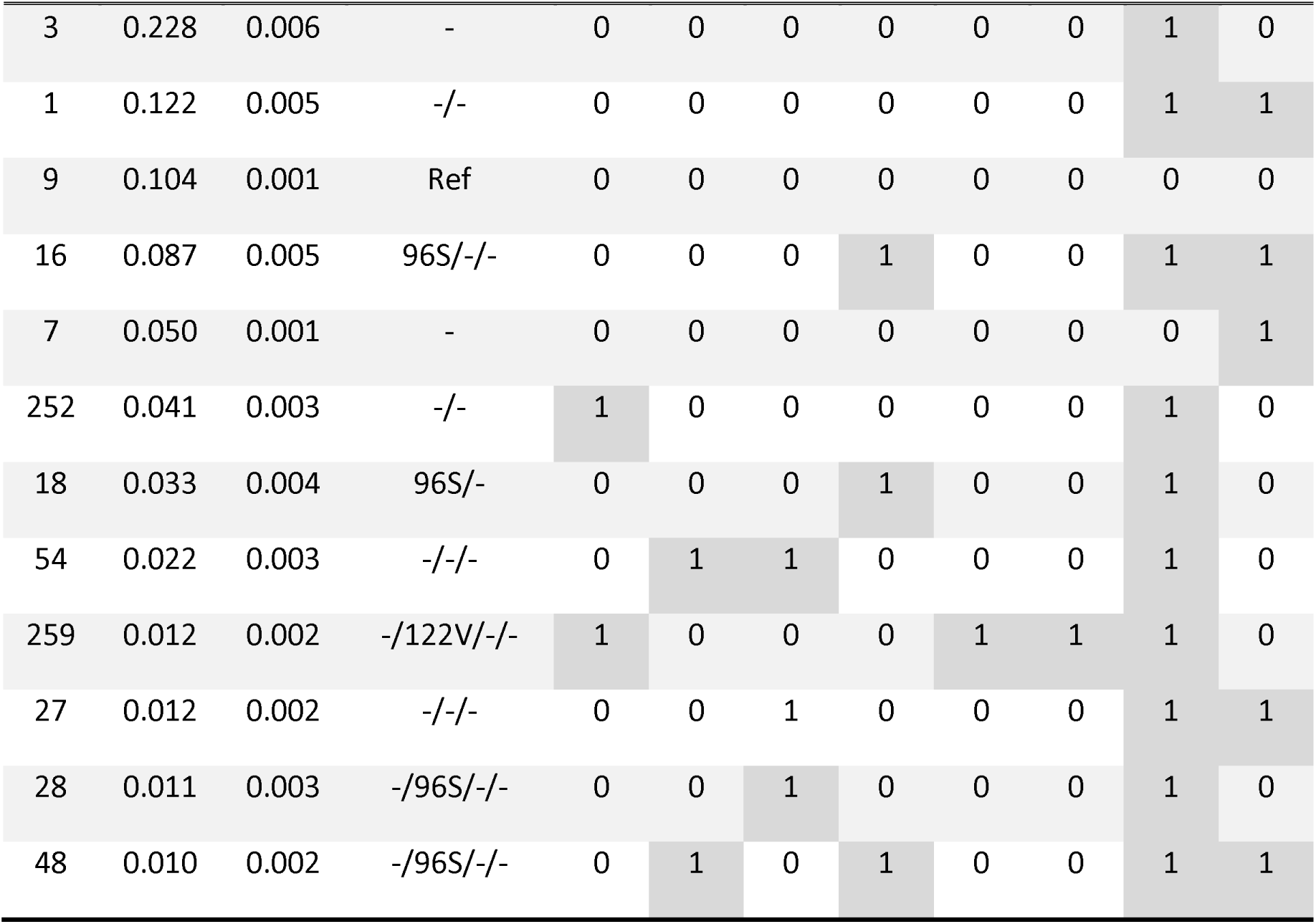
Haplotypes were estimated with PHASE v2.1.1 set to the same parameters in Brandt et al., 2018: Markov chain Monte Carlo (MCMC) samples were taken from a minimum of 100,000 steps, with a discarded burn-in of 10,000; samples were drawn every 100 MCMC steps. Five repetitions were performed, and haplotype frequencies compared to verify consistent assignment. Included are estimates of population haplotypes with frequencies of greater than 1% (count=1262, number haplotypes = 151) and associated estimated standard deviations (S.E.; square root of the variance of the posterior distribution) at 19 variable positions, with 0 representing non-variants and 1 representing variants.

**Table S8.**
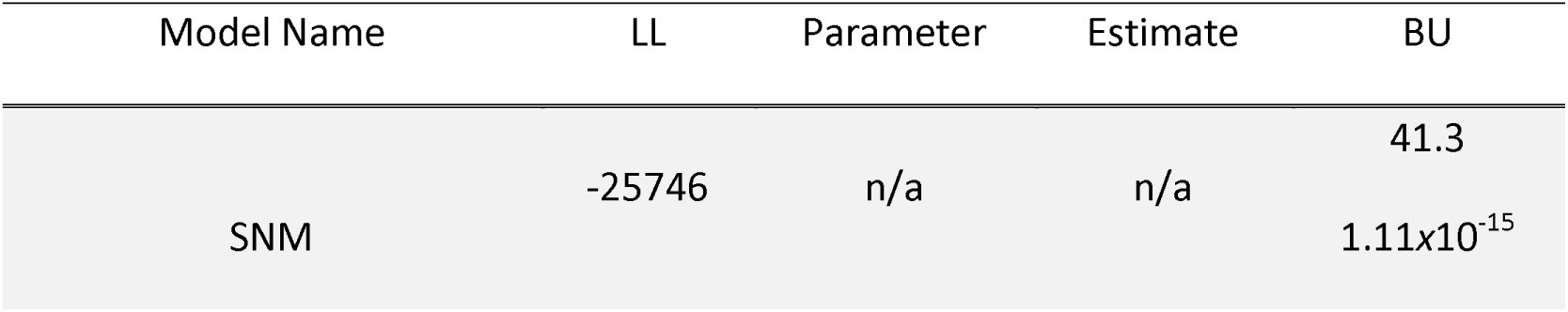
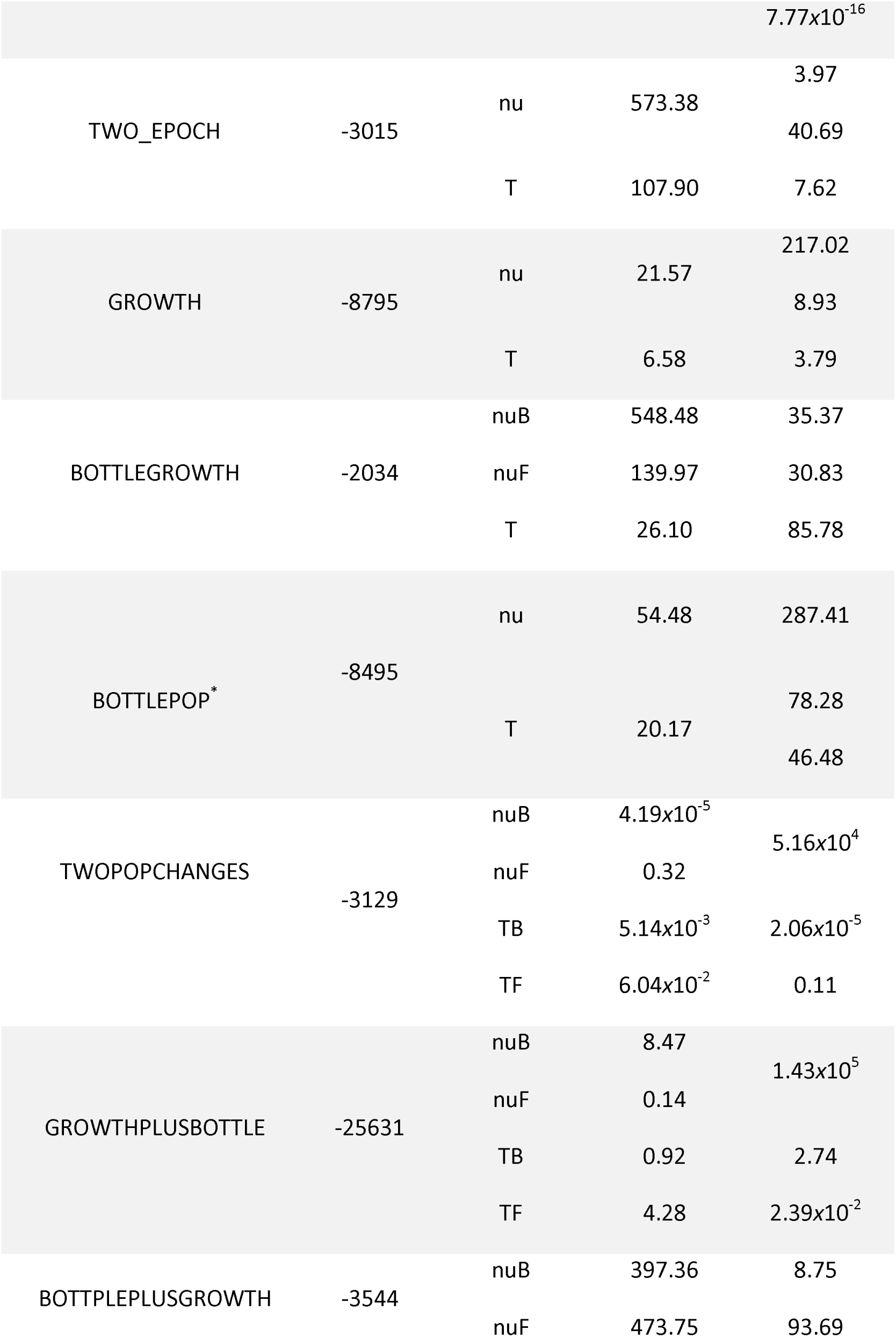
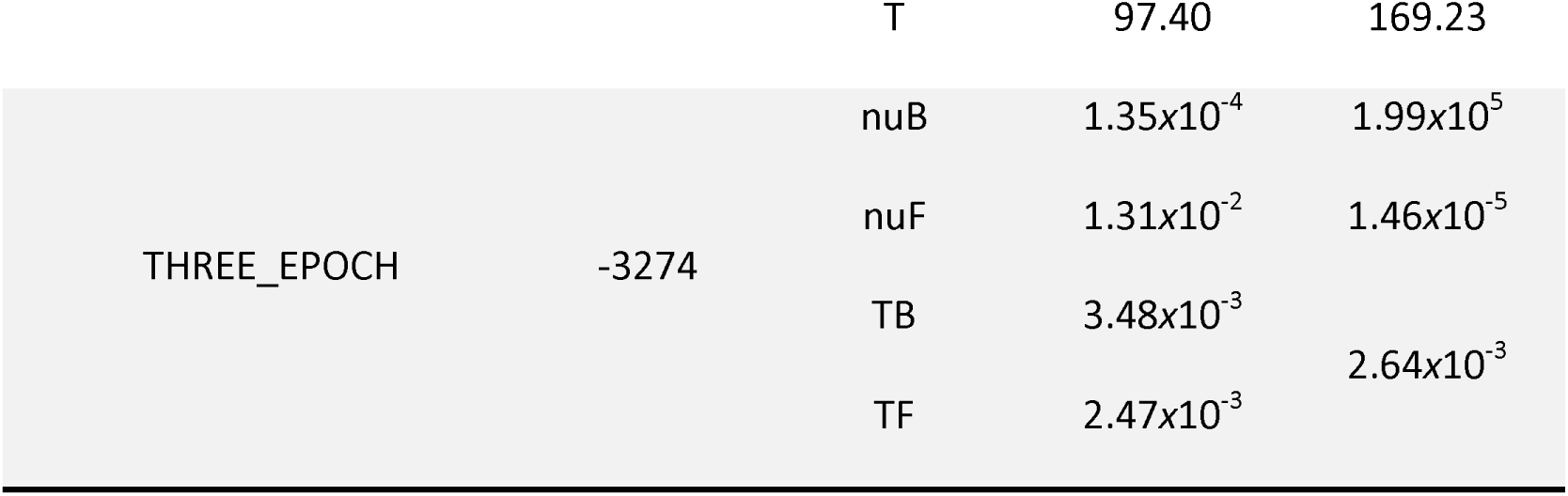
Optimized log-likelihood (LL) and bootstrap uncertainties (BU) obtained from 1D demographic models on Ontario white-tailed deer as a single population. Model specifics and parameters are outlined. The most optimal model for 1D is shown in bold. Modified 1D demographic models are indicated with an asterisk. All time estimates are reported in units of 2*Na generations. Migration rate (m) is reported in units of 2*Na*m.

**Table S9.**
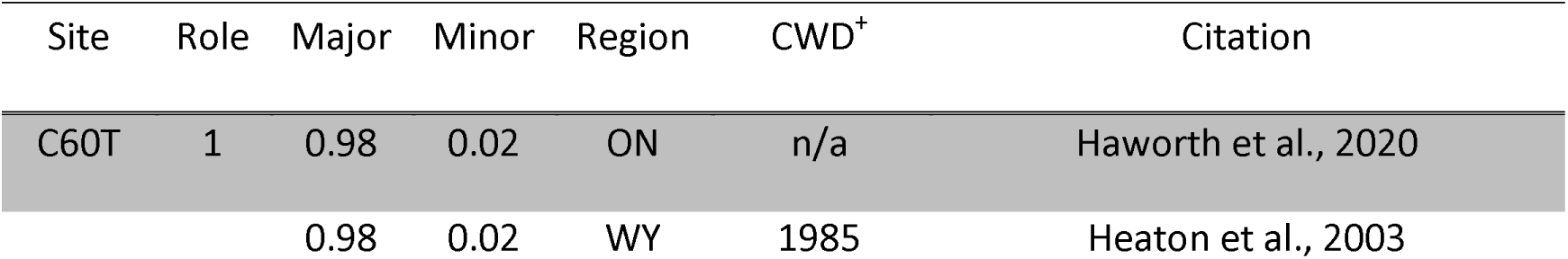
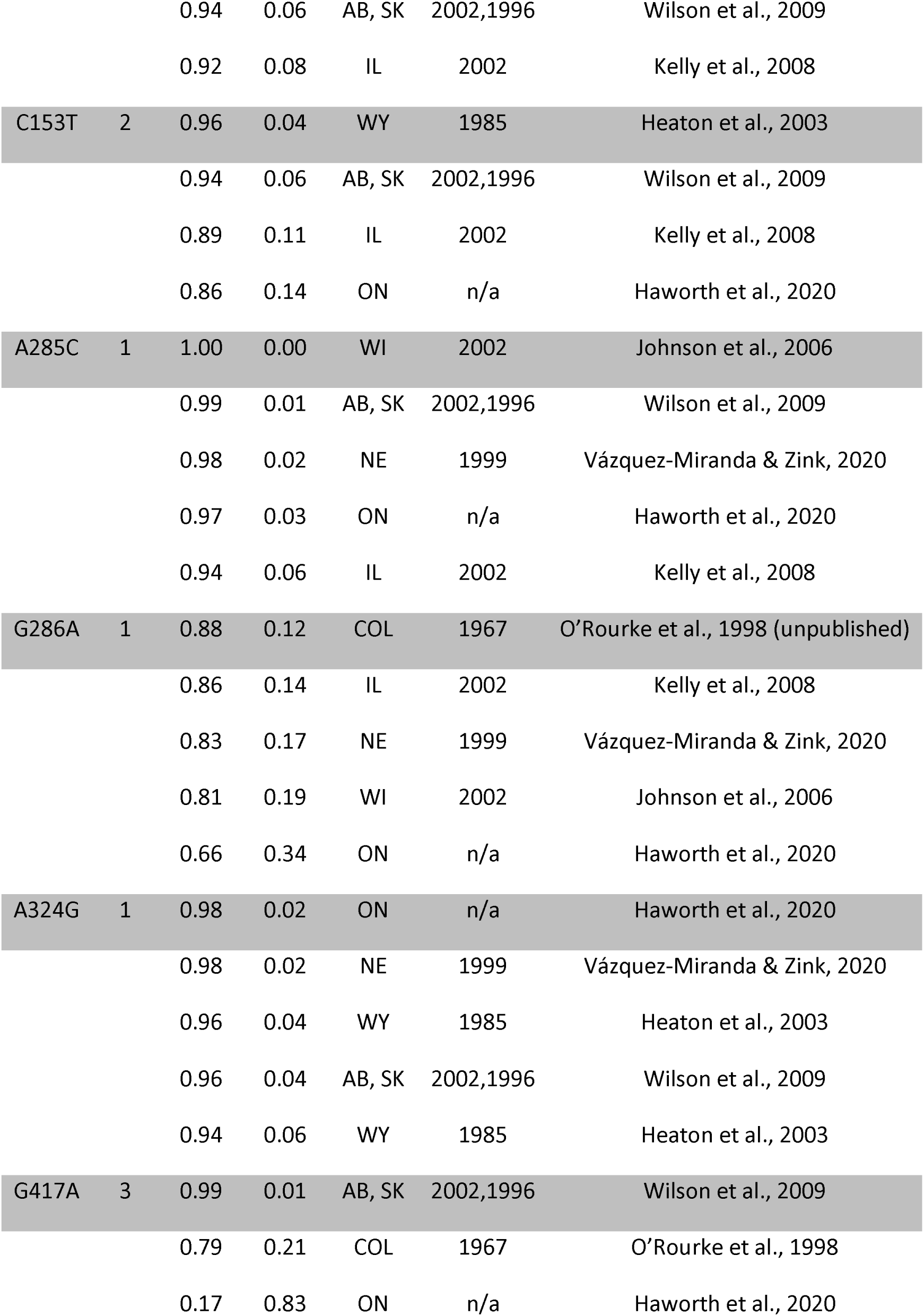
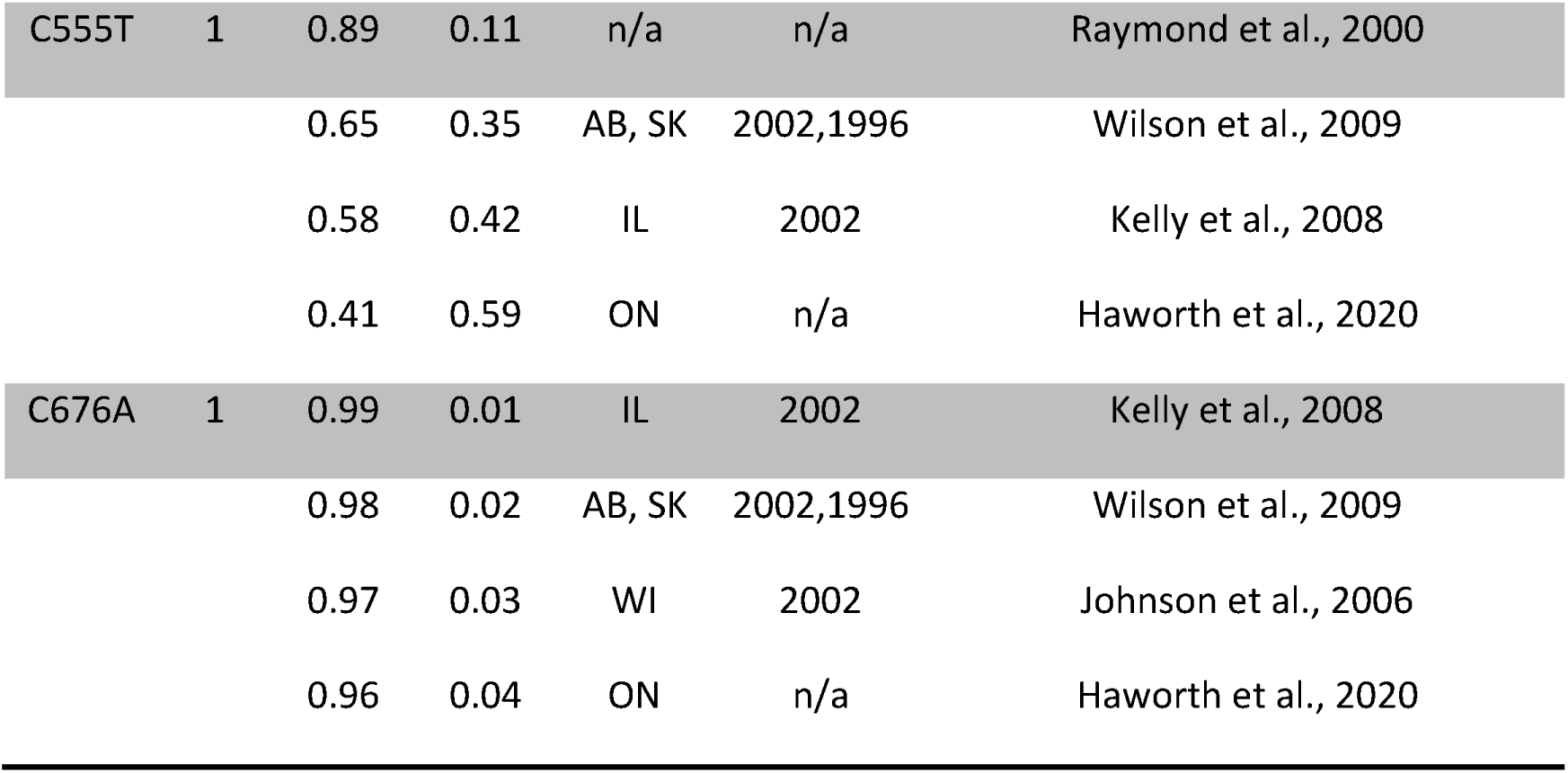
Nucleotide variations in free-ranging white-tailed deer prion protein gene that are associated with chronic wasting disease are either protective (1), increase susceptibility (2), or are neutral. The major and minor allele frequencies for each site across a 771 bp region of the prion protein gene in white-tailed deer are reported for different regions, in descending order. The year CWD was found in free-ranging cervids is reported for each location. The data are from free-ranging white-tailed deer samples collected in: Alberta, Canada (AB); Colorado, USA (COL); Illinois, USA (IL); Ontario, Canada (ON); Nebraska, USA (NE); Saskatchewan, Canada (SK); Wisconsin, USA (WI); and Wyoming, USA (WY).

**Figure S1.**
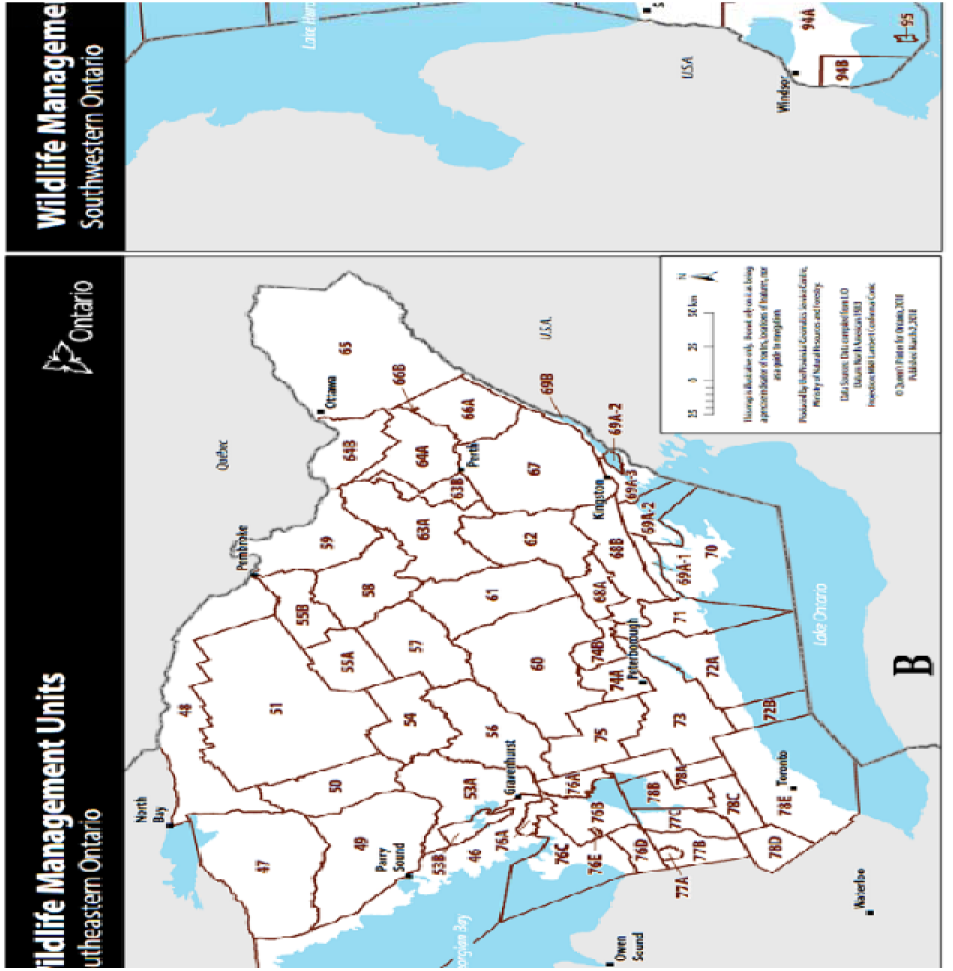
The Canadian province of Ontario as managed by the Ontario Ministry of Natural Resources and Forestry. There are three broad regions Ontario is managed by: (A) Northern Ontario, (B) Southeastern Ontario, and (C) Southwestern Ontario. Collectively (B) and (C) form Southern Ontario. Outlined in red are the wildlife management units designated within each broad region.

**Figure S2.**
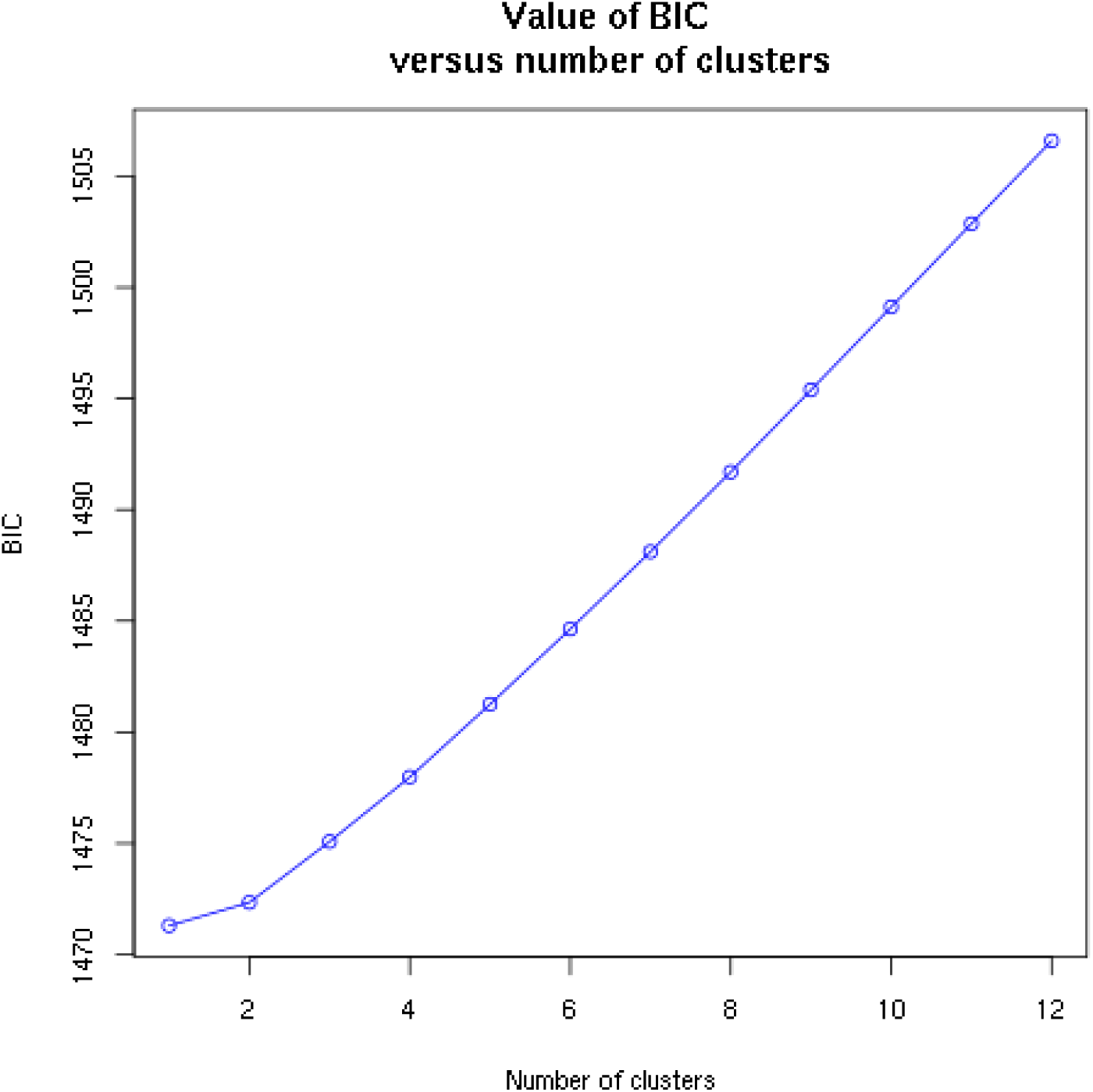
The Bayesian information criterion (BIC) from a population cluster identification using successive K-means cluster assignment on the reduced representation white-tailed deer genome from identified one cluster as optimal.

