## Supplementary Materials for "Characterizing the demographic history and prion protein variation to infer susceptibility to chronic wasting disease in a naïve population of white-tailed deer (*Odocoileus virginianus*)"

**Table S1**. Four degenerate primers used to amplify the functional white-tailed deer prion protein gene region as two overlapping fragments are shown. Fragment 1 is 460 base-pairs in length and Fragment 2 is 590 base-pairs in length.

| Primer | Fragment 1 | Fragment 2 |
| --- | --- | --- |
| Forward (5’-3’) | TCGTCGGCAGCGTCAGATGTGTATAAGAGACAGACRTGGGCATATGATGCTGAYACC | TCGTCGGCAGCGTCAGATGTGTATAAGAGACAGTGGAGGCTGGGGTCAAGG |
| Reverse (5’-3’) | GTCTCGTGGGCTCGGAGATGTGTATAAGAGACAGYTGCCAAAATGTATAAGAGG | GTCTCGTGGGCTCGGAGATGTGTATAAGAGACAGACTACAGGGCTGCAGGTAGAYACT |

**Table S2**. Thermal cycling conditions for PRNP PCR amplification for Fragment 1 and Fragment

| Fragment 1 | Fragment 2 | |
| --- | --- | --- |
| 98℃ for 2 min | | |
| 10 cycles of: | | |
| 98℃ for 15 sec  60-55℃ for 30 sec^1^  72℃ for 45 sec | | 98℃ for 15 sec  65-60℃ for 30 sec^1^  72℃ for 45 sec |
| 98℃ for 15 sec | | |
| 20 cycles of: | | |
| 98℃ for 15 sec  55℃ for 30 sec  72℃ for 45 sec^2^ | | 98℃ for 15 sec  60℃ for 30 sec  72℃ for 45 sec^2^ |
| 72℃ for 7 min | | |
| 1 – decrease by 0.5° per cycle; 2 – increasing by 5 sec per cycle | | |

**Table S3**. Mixture for double restriction enzyme digestion. Samples and reaction mixture were incubated at 37°C for 3 hours and then 25°C overnight on a thermal cycler with a heated lid. Samples and reaction mixture were head-inactivated at 85°C for 30 minutes.

| Reagent | 1 X | Final concentration |
| --- | --- | --- |
| 10x Cutsmart | 2.00 ul | 1 X |
| H_2_O | 11.85 ul |  |
| Msel (10 U/ μl) | 0.10 ul | 1 U |
| Sbfl-HF (20 U/ μl) | 0.05 ul | 1 U |
| Subtotal | 14.00 ul |  |
| DNA | x ul | ~ 500 ng |
| H_2_O | 6.00 – x ul |  |
| Subtotal | 6.00 ul |  |
| Final Volume | 20.00 ul |  |

**Table S4**. Mixture for adapter ligation. The digested fragments were combined with 7 ul of the adapter ligation mixture and 3 ul of a unique barcoded Sbfl adapter (1.0 uM). The ligation on ~ 30 ul reaction mixture was performed in 16°C for 3 hours

| Reagent | 1 X | Final concentration |
| --- | --- | --- |
| 10x CutSmart | 1.00 | 1 X |
| 100 mM ATP | 0.30 | 1 mM |
| H_2_O | 2.45 |  |
| Msel Y adapter (10 uM) | 3.00 | 1 μM |
| T4 DNA ligase (400 U/μl) | 0.25 | 100 U |
| Subtotal | 7.00 ul |  |

**Table S5**. Illumina PCR mixture. The purified restriction-ligation DNA (3 ul) was combined with 7 ul of PCR mixture and a PCR was performed on 10 ul. Four replicated per sample was performed. The thermal cycler profile for this PCR was 98°C for 30 seconds; 20 cycles of 98°C for 20 seconds, 60°C for 30 seconds, 72°C for 45 seconds; and a final extension at 72°C for 5 minutes.

| Reagent | 1 X | Final concentration |
| --- | --- | --- |
| H_2_O | 1.33 |  |
| KAPA HiFi ReadyMix | 5.00 | 2 X |
| PCR primer mix (5 μM each) | 0.67 | 0.5 μM |
| Subtotal | 7.00 ul |  |

**Table S6**. The major/minor allele counts for four nucleotide (nt) positions of variation in the white-tailed deer prion protein gene that are linked to reduced susceptibility or reduced clinical progression of chronic wasting disease for Northern and Southern Ontario are shown. A two-sided Fisher’s Exact test was conducted on the major and minor allele counts at the four chronic wasting disease-linked nucleotide positions. A p-value less than 0.05 was considered significant and indicated that there were significant differences between the groups.

| Location | nt60 | nt285 | nt286 | nt676 |
| --- | --- | --- | --- | --- |
| Northern | 186/3 | 182/7 | 132/57 | 187/2 |
| Southern | 432/10 | 430/12 | 285/157 | 420/22 |
| Southeastern | 306/8 | 308/6 | 196/118 | 300/14 |
| Southwestern | 126/2 | 122/6 | 89/39 | 120/8 |
| p-value | 0.764 | 0.611 | 0.200 | 0.021 |

**Table S7.** Haplotypes were estimated with PHASE v2.1.1 set to the same parameters in Brandt et al., 2018: Markov chain Monte Carlo (MCMC) samples were taken from a minimum of 100,000 steps, with a discarded burn-in of 10,000; samples were drawn every 100 MCMC steps. Five repetitions were performed, and haplotype frequencies compared to verify consistent assignment. Included are estimates of population haplotypes with frequencies of greater than 1% (count=1262, number haplotypes = 151) and associated estimated standard deviations (S.E.; square root of the variance of the posterior distribution) at 19 variable positions, with 0 representing non-variants and 1 representing variants.

| ID | f | S.E. | Codon | 153 | 195 | 198 | 286 | 365 | 378 | 417 | 555 |
| --- | --- | --- | --- | --- | --- | --- | --- | --- | --- | --- | --- |
| 3 | 0.228 | 0.006 | - | 0 | 0 | 0 | 0 | 0 | 0 | 1 | 0 |
| 1 | 0.122 | 0.005 | -/- | 0 | 0 | 0 | 0 | 0 | 0 | 1 | 1 |
| 9 | 0.104 | 0.001 | Ref | 0 | 0 | 0 | 0 | 0 | 0 | 0 | 0 |
| 16 | 0.087 | 0.005 | 96S/-/- | 0 | 0 | 0 | 1 | 0 | 0 | 1 | 1 |
| 7 | 0.050 | 0.001 | - | 0 | 0 | 0 | 0 | 0 | 0 | 0 | 1 |
| 252 | 0.041 | 0.003 | -/- | 1 | 0 | 0 | 0 | 0 | 0 | 1 | 0 |
| 18 | 0.033 | 0.004 | 96S/- | 0 | 0 | 0 | 1 | 0 | 0 | 1 | 0 |
| 54 | 0.022 | 0.003 | -/-/- | 0 | 1 | 1 | 0 | 0 | 0 | 1 | 0 |
| 259 | 0.012 | 0.002 | -/122V/-/- | 1 | 0 | 0 | 0 | 1 | 1 | 1 | 0 |
| 27 | 0.012 | 0.002 | -/-/- | 0 | 0 | 1 | 0 | 0 | 0 | 1 | 1 |
| 28 | 0.011 | 0.003 | -/96S/-/- | 0 | 0 | 1 | 0 | 0 | 0 | 1 | 0 |
| 48 | 0.010 | 0.002 | -/96S/-/- | 0 | 1 | 0 | 1 | 0 | 0 | 1 | 1 |

**Table S8**. Optimized log-likelihood (LL) and bootstrap uncertainties (BU) obtained from 1D demographic models on Ontario white-tailed deer as a single population. Model specifics and parameters are outlined. The most optimal model for 1D is shown in bold. Modified 1D demographic models are indicated with an asterisk. All time estimates are reported in units of 2*Na generations. Migration rate (m) is reported in units of 2*Na*m.

| Model Name | LL | Parameter | Estimate | BU |
| --- | --- | --- | --- | --- |
|  | -25746 | n/a | n/a | 41.3 |
| SNM |  |  |  | 1.11*x*10^-15^ |
|  |  |  |  | 7.77*x*10^-16^ |
|  | -3015 | nu | 573.38 | 3.97 |
| TWO_EPOCH |  |  |  | 40.69 |
|  |  | T | 107.90 | 7.62 |
| GROWTH | -8795 | nu | 21.57 | 217.02 |
|  |  |  |  | 8.93 |
|  |  | T | 6.58 | 3.79 |
| BOTTLEGROWTH | -2034 | nuB | 548.48 | 35.37 |
|  |  | nuF | 139.97 | 30.83 |
|  |  | T | 26.10 | 85.78 |
|  | -8495 | nu | 54.48 | 287.41 |
| BOTTLEPOP^*^ |  | T | 20.17 | 78.28 |
|  |  |  |  | 46.48 |
|  | -3129 | nuB | 4.19*x*10^-5^ | 5.16*x*10^4^ |
| TWOPOPCHANGES |  | nuF | 0.32 |  |
|  |  | TB | 5.14*x*10^-3^ | 2.06*x*10^-5^ |
|  |  | TF | 6.04*x*10^-2^ | 0.11 |
| GROWTHPLUSBOTTLE | -25631 | nuB | 8.47 | 1.43*x*10^5^ |
|  |  | nuF | 0.14 |  |
|  |  | TB | 0.92 | 2.74  2.39*x*10^-2^ |
|  |  | TF | 4.28 |  |
| BOTTPLEPLUSGROWTH | -3544 | nuB | 397.36 | 8.75 |
|  |  | nuF | 473.75 | 93.69 |
|  |  | T | 97.40 | 169.23 |
| THREE_EPOCH | -3274 | nuB | 1.35*x*10^-4^ | 1.99*x*10^5^ |
|  |  | nuF | 1.31*x*10^-2^ | 1.46*x*10^-5^ |
|  |  | TB | 3.48*x*10^-3^ | 2.64*x*10^-3^ |
|  |  | TF | 2.47*x*10^-3^ |  |

**Table S9**. Nucleotide variations in free-ranging white-tailed deer prion protein gene that are associated with chronic wasting disease are either protective (1), increase susceptibility (2), or are neutral. The major and minor allele frequencies for each site across a 771 bp region of the prion protein gene in white-tailed deer are reported for different regions, in descending order. The year CWD was found in free-ranging cervids is reported for each location. The data are from free-ranging white-tailed deer samples collected in: Alberta, Canada (AB); Colorado, USA (COL); Illinois, USA (IL); Ontario, Canada (ON); Nebraska, USA (NE); Saskatchewan, Canada (SK); Wisconsin, USA (WI); and Wyoming, USA (WY).

| Site | Role | Major | Minor | Region | CWD^+^ | Citation |
| --- | --- | --- | --- | --- | --- | --- |
| C60T | 1 | 0.98 | 0.02 | ON | n/a | Haworth et al., 2020 |
|  |  | 0.98 | 0.02 | WY | 1985 | Heaton et al., 2003 |
|  |  | 0.94 | 0.06 | AB, SK | 2002,1996 | Wilson et al., 2009 |
|  |  | 0.92 | 0.08 | IL | 2002 | Kelly et al., 2008 |
| C153T | 2 | 0.96 | 0.04 | WY | 1985 | Heaton et al., 2003 |
|  |  | 0.94 | 0.06 | AB, SK | 2002,1996 | Wilson et al., 2009 |
|  |  | 0.89 | 0.11 | IL | 2002 | Kelly et al., 2008 |
|  |  | 0.86 | 0.14 | ON | n/a | Haworth et al., 2020 |
| A285C | 1 | 1.00 | 0.00 | WI | 2002 | Johnson et al., 2006 |
|  |  | 0.99 | 0.01 | AB, SK | 2002,1996 | Wilson et al., 2009 |
|  |  | 0.98 | 0.02 | NE | 1999 | Vázquez-Miranda & Zink, 2020 |
|  |  | 0.97 | 0.03 | ON | n/a | Haworth et al., 2020 |
|  |  | 0.94 | 0.06 | IL | 2002 | Kelly et al., 2008 |
| G286A | 1 | 0.88 | 0.12 | COL | 1967 | O’Rourke et al., 1998 (unpublished) |
|  |  | 0.86 | 0.14 | IL | 2002 | Kelly et al., 2008 |
|  |  | 0.83 | 0.17 | NE | 1999 | Vázquez-Miranda & Zink, 2020 |
|  |  | 0.81 | 0.19 | WI | 2002 | Johnson et al., 2006 |
|  |  | 0.66 | 0.34 | ON | n/a | Haworth et al., 2020 |
| A324G | 1 | 0.98 | 0.02 | ON | n/a | Haworth et al., 2020 |
|  |  | 0.98 | 0.02 | NE | 1999 | Vázquez-Miranda & Zink, 2020 |
|  |  | 0.96 | 0.04 | WY | 1985 | Heaton et al., 2003 |
|  |  | 0.96 | 0.04 | AB, SK | 2002,1996 | Wilson et al., 2009 |
|  |  | 0.94 | 0.06 | WY | 1985 | Heaton et al., 2003 |
| G417A | 3 | 0.99 | 0.01 | AB, SK | 2002,1996 | Wilson et al., 2009 |
|  |  | 0.79 | 0.21 | COL | 1967 | O’Rourke et al., 1998 |
|  |  | 0.17 | 0.83 | ON | n/a | Haworth et al., 2020 |
| C555T | 1 | 0.89 | 0.11 | n/a | n/a | Raymond et al., 2000 |
|  |  | 0.65 | 0.35 | AB, SK | 2002,1996 | Wilson et al., 2009 |
|  |  | 0.58 | 0.42 | IL | 2002 | Kelly et al., 2008 |
|  |  | 0.41 | 0.59 | ON | n/a | Haworth et al., 2020 |
| C676A | 1 | 0.99 | 0.01 | IL | 2002 | Kelly et al., 2008 |
|  |  | 0.98 | 0.02 | AB, SK | 2002,1996 | Wilson et al., 2009 |
|  |  | 0.97 | 0.03 | WI | 2002 | Johnson et al., 2006 |
|  |  | 0.96 | 0.04 | ON | n/a | Haworth et al., 2020 |

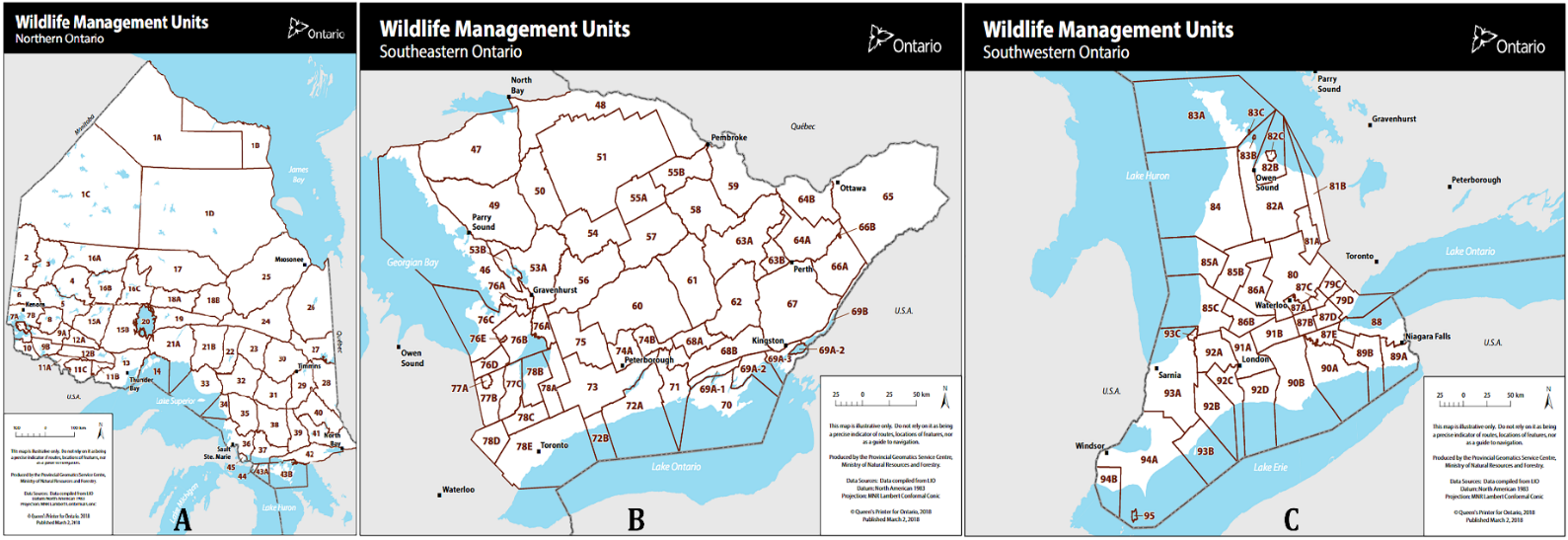

**Figure S1**. The Canadian province of Ontario as managed by the Ontario Ministry of Natural Resources and Forestry. There are three broad regions Ontario is managed by: (A) Northern Ontario, (B) Southeastern Ontario, and (C) Southwestern Ontario. Collectively (B) and (C) form Southern Ontario. Outlined in red are the wildlife management units designated within each broad region.

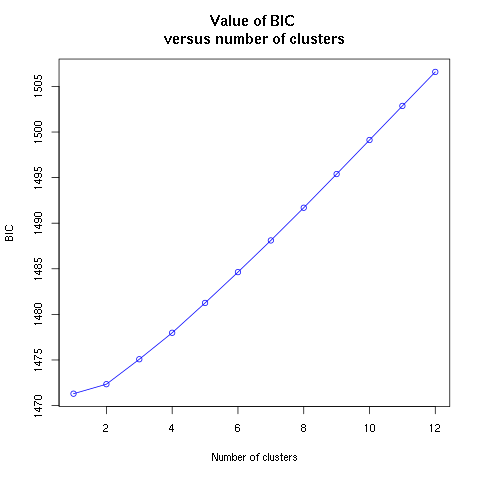

**Figure S2**. The figure shows the value of the Bayesian information criterion (BIC) versus the number of clusters analyzed. The BIC from a population cluster identification using successive K-means cluster assignment on the reduced representation white-tailed deer genome from identified one cluster as optimal.
